# The capacity of the medial temporal lobe to represent memory items in their ordinal position in a sequence is domain-general

**DOI:** 10.1101/2024.09.25.614993

**Authors:** Ainsley Temudo, Nina Dolfen, Bradley R. King, Genevieve Albouy

## Abstract

Memory systems in humans are less segregated than initially thought as learning tasks from different memory domains (e.g., declarative vs. procedural) can recruit similar brain areas. However, it remains unclear whether the functional role of these overlapping brain regions – and the hippocampus in particular - is domain-general. Here, we test the hypothesis that the hippocampus encodes and preserves the temporal order of sequential information irrespective of the nature of that information. We used multivariate pattern analyses (MVPA) of functional magnetic resonance imaging (fMRI) data acquired in young healthy individuals during the execution of learned sequences of movements and objects to test whether the hippocampus represents information about the temporal order of items in a learned sequence irrespective of their nature. We also examined such coding in brain regions involved in both motor (primary and premotor cortices) and object (perirhinal cortex and parahippocampus) sequence learning. Our results suggest that hippocampal and perirhinal multivoxel activation patterns do not carry information about specific items or temporal position in a random series of objects or movements. Rather, these regions code for the representation of items in their learned temporal position in sequences irrespective of their nature (i.e., item-position coding). In contrast, although all other ROIs showed evidence of item-position coding, this representation could - at least partially - be attributed to the coding of other information such as position information. Altogether, our findings indicate that the capacity of regions in the medial temporal lobe to represent the temporal order of sequential information is domain-general. Our data suggest that these regions contribute to the development of item-position maps that might provide a cognitive framework to order sequential behaviors irrespective of their nature.

## Introduction

Influential models of memory organization suggest that the declarative and procedural memory systems are supported by distinct brain networks in humans [1]. This framework was largely based on neuropsychological studies showing that patients with hippocampal lesions exhibited severe declarative learning deficits but could learn new procedural (motor) skills [2,3]. However, more recent evidence suggests that these two memory systems are less segregated than initially thought [4,5]. For example, a large number of neuroimaging studies indicate that the hippocampus is involved in motor sequence learning. (e.g., [6–12]) and that these hippocampal responses are related to the subsequent memory consolidation process [7,9]. This is in line with more recent studies in patients with hippocampal lesions demonstrating that the hippocampus is necessary for consolidation even when it is not required for initial learning [13,14]. Despite the evidence reviewed above suggesting that different memory systems can recruit common brain areas, it remains unclear whether the functional role of these overlapping brain regions – in particular the hippocampus – is shared across memory domains.

Indirect evidence for potential overlapping hippocampal function between memory domains comes from studies using multivariate analyses of functional Magnetic Resonance Imaging (fMRI) data. This research suggests that the hippocampus plays a critical role in the processing of temporal information in both the declarative (e.g., [15]) and the procedural memory domain [16]. In the declarative domain, results show that hippocampal activity patterns contain information about the temporal order of episodic events [17] as well as items in learned sequences of letters [18] and objects [15]. Recent work from our group has extended these findings to the procedural memory domain and has shown that hippocampal activation patterns carry information about the position of finger movements in a learned motor sequence [16]. These findings are in line with a recent influential model of hippocampal function, i.e. the *hippocampal sequencing hypothesis*, which proposes that the hippocampus produces a “*sequential content-free structure*” to organize experiences distributed across different cortical nodes that might represent *content-specific* representations [19]. This theory conceptualizes the hippocampus as “*a general-purpose sequence generator that carries content-limited ordinal structure, and tiles the gaps between events or places to be linked*” [20]. Based on this theory and the evidence reviewed above, it is tempting to speculate that the capacity of the hippocampus to encode the order of learned information does not depend on the nature of the memory items to be ordered. This, however, remains hypothetical as there is no empirical evidence supporting such view.

The aim of this study was therefore to empirically test this hypothesis. To do so, we designed a serial reaction time (SRT) task that allowed us to probe sequence learning in both the motor and the declarative memory domains (i.e., memorizing a sequence of finger movements and a sequence of objects, respectively). We used multivoxel representational similarity analysis (RSA) of fMRI data acquired during sequence task performance to examine whether hippocampal activation patterns carry information about the order of items in a learned sequence irrespective of the memory domain. Based on the evidence presented above, we hypothesized that hippocampal activation patterns would represent information about the temporal order of memory items in a sequence within each memory domain (i.e., separately for movements [16] and objects [15]), but also regardless of the memory domain (i.e., irrespective of whether the item in a particular position is a movement or an object). We also examined such coding in brain regions involved in motor (primary motor and premotor cortices, [16]) and object (perirhinal cortex and parahippocampus, [15]) sequence learning. We expected item (finger / object) and position information (temporal position in the task stream) to be coded in memory domain-specific regions. Specifically, we hypothesized that fingers and objects would be coded in M1 and in the perirhinal cortex, respectively and that position information would be coded in the premotor and parahippocampal cortices [15–16].

## Results

Participants performed a serial reaction time task (SRTT, Fig. 1A) and learned on the first experimental day a sequence of finger movements (motor sequence, SQ MOT) and a sequence of objects (object sequence, SQ OBJ) during two separate experimental sessions the order of which was counterbalanced across individuals (Fig. 1B). On day 2 of the experiment, participants’ brain activity was recorded using fMRI while they practiced the learned motor and object sequences. They also practiced a random (RD SRTT) control task involving random movements and random object presentation at the start of each experimental session on day 1 to measure baseline performance and inside the scanner to provide a control condition for the RSA (Fig. 1A-B). Performance on the three task conditions was retested outside the scanner to probe memory retention (Retest, Fig. 1B).

**Figure 1.**
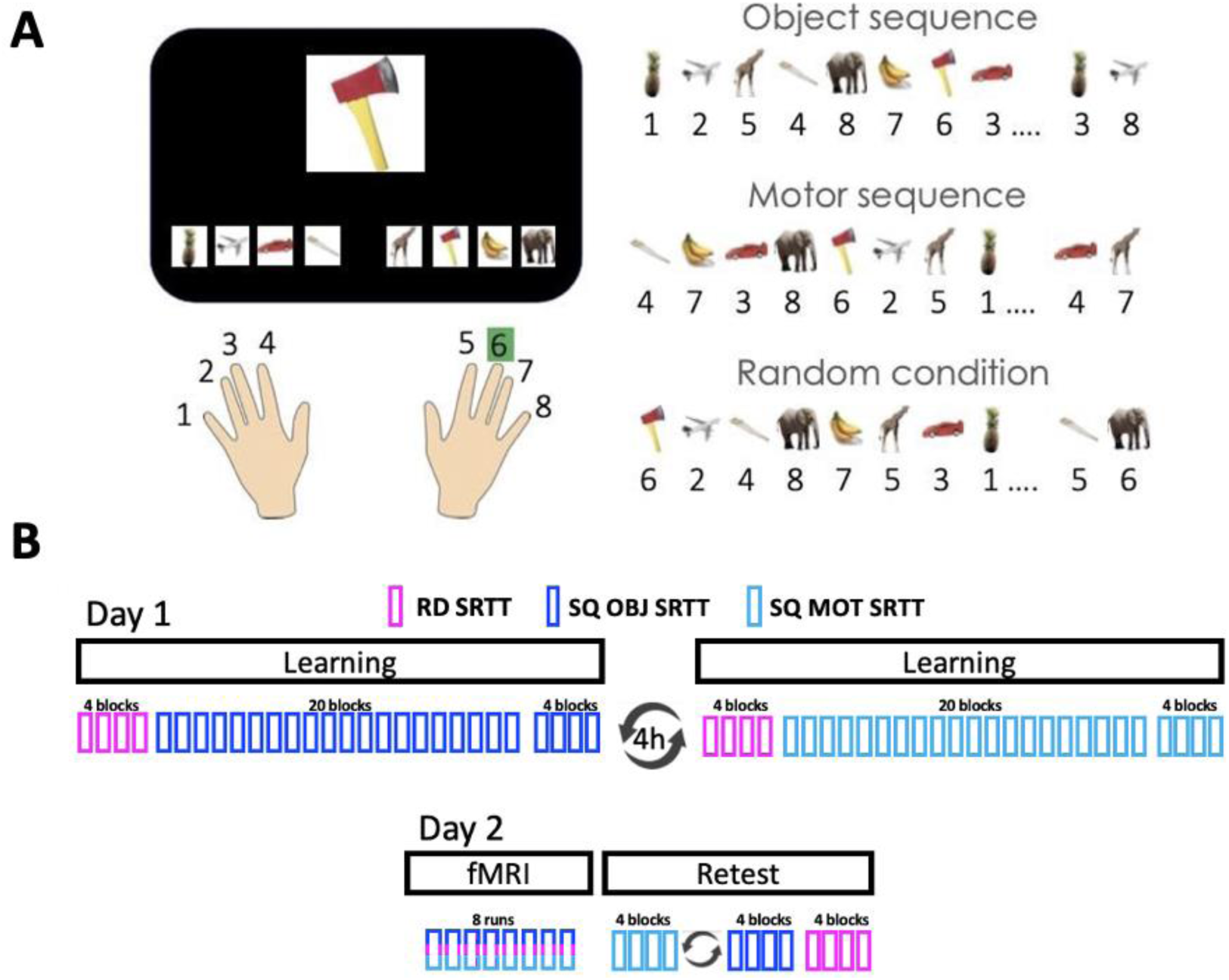
**A.** *Serial reaction time task (SRTT*). Left panel. A stimulus appears in the center of the screen and participants are instructed to respond as fast and as accurately as possible according to the finger/object mapping displayed at the bottom of the screen. Note that the finger-object mapping displayed on the bottom of the screen changed after each stream of 8 elements (i.e., every 8 objects / 8 finger presses) in order to orthogonalize object and motor sequence learning (see right panel). Right panel. Three different task conditions were designed: object sequence with random finger presses (that resulted in the learning of an 8-element object sequence, SQ OBJ), motor sequence with random object presentation (that resulted in the learning of an 8-element finger sequence, SQ MOT) and a random series with random finger presses and random object presentation (RD). Numbers in the figure represent fingers whereby 1 and 8 correspond to the left and right little fingers, respectively. **B.** *Experimental Design*. Top panel. On day 1, after baseline performance assessment with a random (RD) SRTT, participants learned a sequence of objects (SQ OBJ SRTT) and a sequence of movements (SQ MOT SRTT), outside the MRI scanner, in two separate sessions 4h apart (order counterbalanced across participants). Bottom panel. On day 2, brain activity was recorded on the three task conditions with fMRI over 8 runs during which the RD, SQ OBJ and SQ MOT conditions were pseudo-randomly interleaved so that the same condition did not repeat more than 2 times in a row. A retest session was then performed outside the scanner to assess memory retention on each sequence condition (order counterbalanced across participants) followed by a final test on the random condition.

### Behavioral results

Analyses of baseline performance extracted from the random SRTT acquired at the start of each learning session (Fig. 1) are reported in section 1 of the supplementary results. Analyses of the sequence SRTT data indicated that performance speed (i.e., mean response time) improved during learning on day 1 for both the motor and object sequence tasks (block effect; training: F(19,551)=38.98, ɳ_p_^2^=0.57, p<0.001; test: F(3,87)=3.78, ɳ_p_^2^=0.12, p=0.01) but the motor task presented overall faster performance (condition by block effect; training: F(19,551)=2.72, ɳ_p_^2^= 0.09, p<0.001; test: F(3,87)=1.57, ɳ_p_^2^=0.05, p=0.2; condition effect; training: F(1,29)=5.65, ɳ_p_^2^=0.16, p=0.02; test: F(1,29)=9.45, ɳ_p_^2^= 0.25, p=0.005; Fig. 2, day 1, top panel). In contrast, performance accuracy remained stable during learning on day 1 and similar between conditions (training: condition by block effect: F(19,551)=0.61, ɳ_p_^2^=0.02, p=0.9; block effect: F(19,551)=1.43, ɳ_p_^2^=0.05, p=0.11; condition effect: F(1,29)=3.25, ɳ_p_^2^=0.1, p=0.08; test: condition by block effect: F(3,87)=2.84, ɳ_p_^2^=0.09, p=0.04; block effect F(3,87)=0.72, ɳ_p_^2^=0.02, p=0.55; condition effect: F(1,29)=1.2, ɳ_p_^2^=0.04, p=0.28; Fig. 2, day 1, bottom panel).

**Figure 2.**
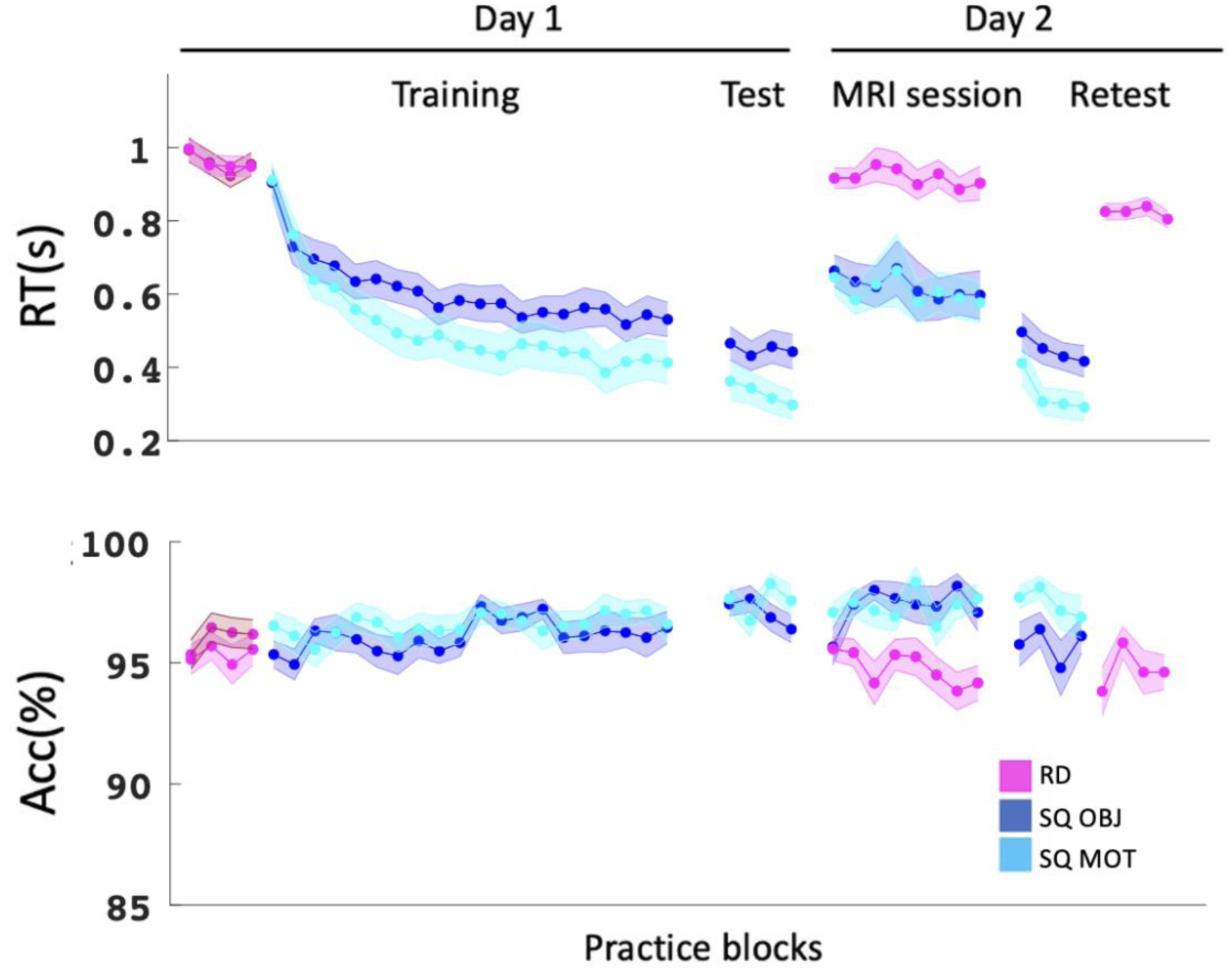
Behavioral results. Response time in seconds (top panel) and accuracy (% correct response, bottom panel) across blocks and sessions for all task conditions. Dark blue: object sequence (SQ OBJ), light blue: motor sequence (SQ MOT), pink: random blocks (RD) (darker shade corresponds to random practice during the object session). Shaded error bars=SEM.

Data collected on day 2 in the MRI scanner indicated that performance remained stable across runs (block effect; speed: F(7,203)=0.92, ɳ_p_^2^=0.03, p=0.5; accuracy: block effect: F(7,203)=1.23, ɳ_p_^2^= 0.04, p=0.29). Importantly, performance was better for the sequence conditions as compared to random condition (condition effect; speed: F(2,58)=66.39, ɳ_p_^2^=0.7, p<0.001; motor vs. random: p<0.001; object vs. random: p<0.001; accuracy: F(2,58)=26.28, ɳ_p_^2^=0.48, p<0.001; motor vs. random: p<0.001; object vs. random: p<0.001) but similar between sequence conditions (speed and accuracy: object vs. motor: ps>0.99; Fig. 2, MRI session). During the retest performed outside the scanner, performance speed improved across practice blocks (block effect: F(3,87)=8.52, ɳ_p_^2^=0.23, p<0.001) while accuracy remained stable (block effect: F(3,87)=2.8, ɳ_p_^2^=0.09, p=0.05). Importantly, performance was overall better for sequence conditions when compared to random (condition effect; speed: F(2,58)=126.8; ɳ_p_^2^=0.81, p<0.001; motor vs. random: p<0.001; object vs. random: p<0.001; accuracy: condition effect; F(2,58)=8.31, ɳ_p_^2^=0.22, p <0.001; motor vs. random: p<0.002; object vs. random: p=0.31) and performance for the motor task remained better than for the object task (motor vs. object; speed: p=0.001; accuracy: p=0.08; Fig. 2, Retest).

Overall, these results suggest that participants specifically learned and retained both the motor and object sequences, although to a different extent as performance was better on the motor as compared to the object task (see discussion and section 3 of the supplemental results for brain-behavior correlation analyses). Importantly, performance on the two sequence tasks was similar in the scanner and a sequence-specific performance advantage was observed despite the interleaved nature of practice.

### fMRI results

We performed representational similarity analyses (RSA) of the fMRI data collected while participants were performing the sequence learning tasks on day 2 (see Fig. 1B). These analyses are based on the assumption that if a particular type of information is processed by a particular region of interest (ROI), then it will be represented in the pattern of activation across voxels in that brain region [21]. Such analyses therefore predict higher correlations in voxel activity between pairs of trials sharing this information as compared to pairs of trials not sharing this information (see diagonal and off-diagonal cells in exemplary matrices depicted in Fig. 3A, 4A and 5A). This approach was used to test whether our ROIs represent information about specific items (fingers or objects) in their learned temporal position in the sequence within each memory domain (i.e., finger-position coding in the motor domain and object-position coding the declarative memory domain) but also regardless of the memory domain (i.e., item-position coding across domains). We also used the random task data to test whether the ROIs carried information about (1) items irrespective of their position in the pattern (i.e., finger / object coding), and (2) temporal positions irrespective of item (i.e., position coding). Analyses were performed on five bilateral ROIs involved in sequence learning [15,16]: the primary motor cortex (M1) and premotor cortex (PMC) for motor sequence learning; the perirhinal cortex (PER) and parahippocampus (PHC) for object sequence learning; and the hippocampus (HC) for both motor and object sequence learning tasks. Results are presented below within each memory domain first and between memory domains last. Correction for multiple comparisons was performed using FDR correction for 5 ROIs in both the within- and between-memory domains analyses.

**Figure 3.**
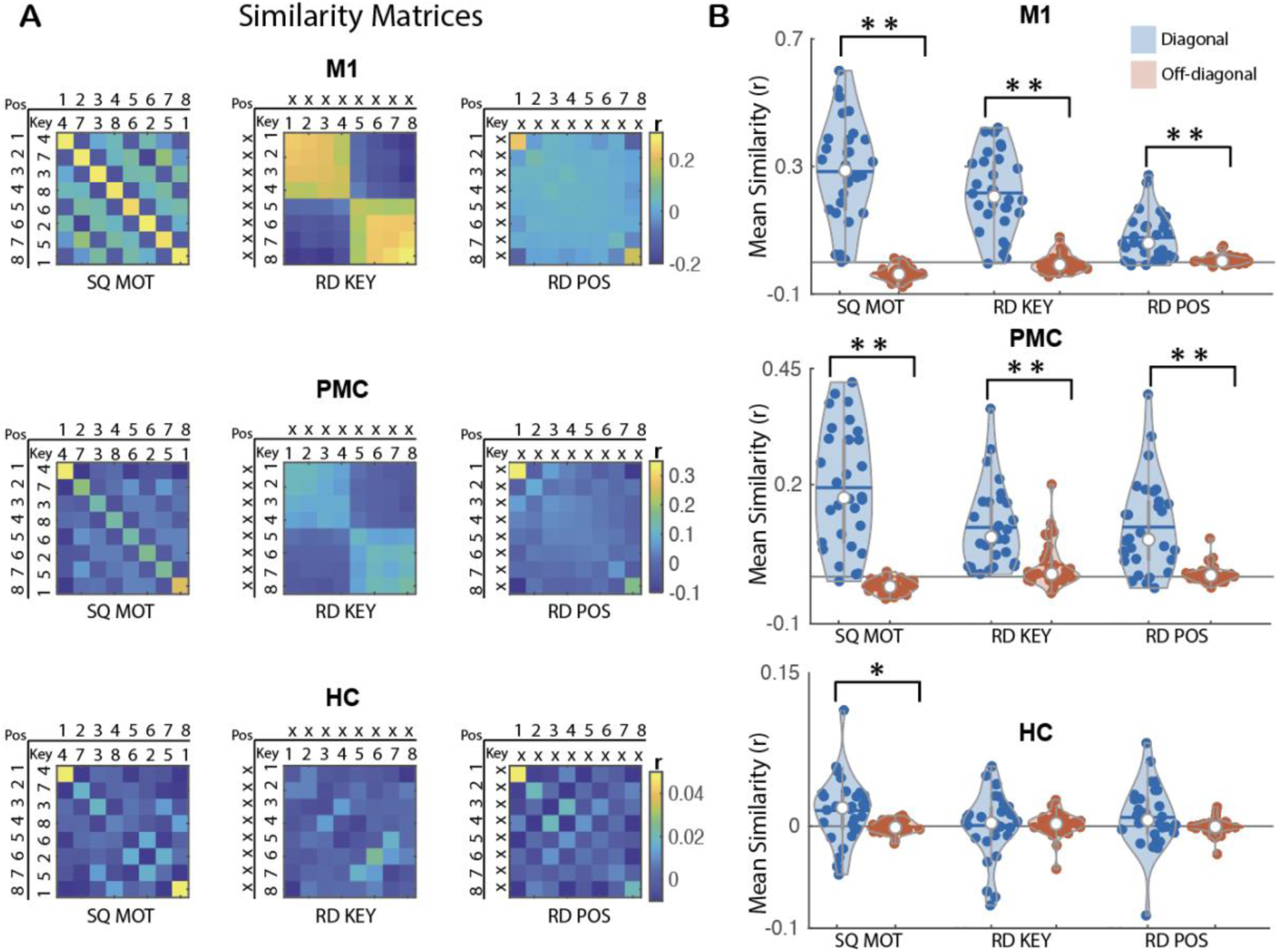
MVPA results for motor sequence learning. **A**. Group average neural similarity matrices for the ROIs involved in motor learning. Pattern similarity was computed across repetitions of the motor sequence to assess finger-position coding in the sequence condition (SQ MOT matrix, left panel) as well as across repetitions of the random patterns to quantify finger/key (RD KEY matrix, middle panel) and position (RD POS matrix, right panel) coding in the random condition. Color bar represents mean similarity (r). In the POS rows/columns, numbers represent the temporal position in the 8-element sequence or in the random stream. In the KEY rows/columns, numbers represent fingers. (X) represents a random position or key in key and position matrices, respectively. **B**. Mean pattern similarity for diagonal (blue) and off-diagonal (red) cells as a function of matrices and ROIs. Results indicate that M1 and PMC show evidence of finger-position coding in the sequence condition as well as finger and position coding during random practice. In contrast, the hippocampus (HC) shows evidence for finger-position coding in the sequence condition but not for finger or position coding in the random condition. Asterisks indicate significant differences between diagonal and off-diagonal (one sided paired sample t-test; Bonferroni corrected \**p_corr_*<.05 and \*\**p_corr_*<.001). Colored circles represent individual data, jittered in arbitrary distances on the x-axis to increase perceptibility. Horizontal lines represent means and white circles represent medians. The shape of the violin [22] depicts the kernel density estimate of the data. Note that as in earlier research [16], Y axis scales are different between ROIs to accommodate for differences in signal-to-noise ratio (and therefore in effect sizes) between ROIs.

### Motor memory domain

Within-motor memory domain analyses first examined *selective coding* in the ROIs. This was assessed by comparing the mean correlation of the diagonal elements of the representational similarity matrices with the mean correlation of the off-diagonal elements (see Fig. 3A). Specifically, we measured the extent to which specific ROIs represent information about (1) fingers in their learned temporal position in the sequence (i.e., finger-position coding in the sequence condition), (2) fingers irrespective of their position (i.e., finger coding in the random condition), and (3) temporal positions irrespective of the finger (i.e., position coding in the random condition). Next, when the ROIs presented evidence of coding for multiple representations (e.g., both finger-position and finger coding), we performed linear regression analyses to assess the *contribution of the different representations* to finger-position coding. Within-motor memory domain analyses were performed on all 5 ROIs, but only the results corresponding to the 3 ROIs involved in motor sequence learning (i.e., the primary motor cortex M1, the premotor cortex PMC and the hippocampus HC; [16] are presented in the main text (see supplemental Figures S1 and S2 and Table S4 for results related to the other ROIs).

#### Finger-position coding in a sequence

To investigate whether the motor sequence ROIs represent information about fingers and their learned temporal position in the sequence, we computed the correlation in voxel activity patterns between pairs of individual key presses across repetitions of the motor sequence. This resulted in an 8×8 motor sequence condition neural similarity matrix for each motor ROI (Fig. 3A). Analysis of this matrix (***SQ MOT***) revealed significantly higher mean similarity along the diagonal (i.e., same finger + position) as compared to the off-diagonal (i.e., different finger + position) in all motor ROIs (paired sample t-test: M1, t(29)=9.72, d=1.78, p_corr_<0.005; PMC, t(29)=8.2, d=1.5, p_corr_<0.005; HC, t(29)=2.47, d=0.45, p_corr_=0.05, Fig. 3B). These results indicate that all motor ROIs carry information about fingers in their learned temporal position in the sequence.

#### Finger and position coding in the random condition

As each finger movement in the sequence condition was always executed at the same temporal position, any similarity effects observed in the finger-position analysis described above could be driven by the coding of finger and/or position information. Control analyses were therefore performed using the random data to test whether the motor ROIs also represented information about finger and position. To assess finger coding, we computed similarity in activation patterns between individual key presses across repetitions of the random series. The resulting 8×8 matri*x* (***RD KEY***, Fig. 3A) revealed significantly higher mean similarity along the diagonal (i.e., same finger/key) as compared to the off-diagonal (i.e., different finger/key) in M1 and PMC (paired sample t-test: M1, t(29)=9.95, d=1.82, p_corr_<0.005; PMC, t(29)=7.17, d=1.31, p_corr_<0.005) but not in the HC (t(29)=-0.19, d=-0.04, p_corr_=1, Fig. 3B). To assess position coding irrespective of the finger, we computed similarity in activation patterns between individual positions in a sequence across repetitions of the random series. The resulting 8×8 matrix (***RD POS***, Fig. 3A) showed significantly higher mean similarity along the diagonal (i.e., same position) as compared to the off-diagonal (i.e., different position) in M1 and PMC (paired sample t-test: M1, t(29)=5.74, d=1.05, p_corr_<0.005; PMC, t(29)=6.12, d=1.12, p_corr_<0.005) but not in the HC (t(29)=1.45, d=0.26, p_corr_=0.4, Fig. 4B). Importantly, similar to our earlier research [16], position coding was particularly pronounced at the edges of the stream of 8 elements (i.e., start and end of the stream – see top left and bottom right corner cells in *RD POS* matrix in Fig. 3A). To verify whether position coding was driven by these boundary effects, we performed control analyses excluding boundary positions from the position matrix. These control analyses showed that position coding only remained significant in the PMC (see supplementary Table S1 for corresponding statistics) and therefore suggest that PMC carries information about position (independent of the strong boundary effects) under random conditions while position coding in M1 appears to be heavily influenced by boundary information. Altogether, these results indicate that M1 and PMC carry information about finger and position (PMC) or boundary position (M1) during random practice.

**Figure 4.**
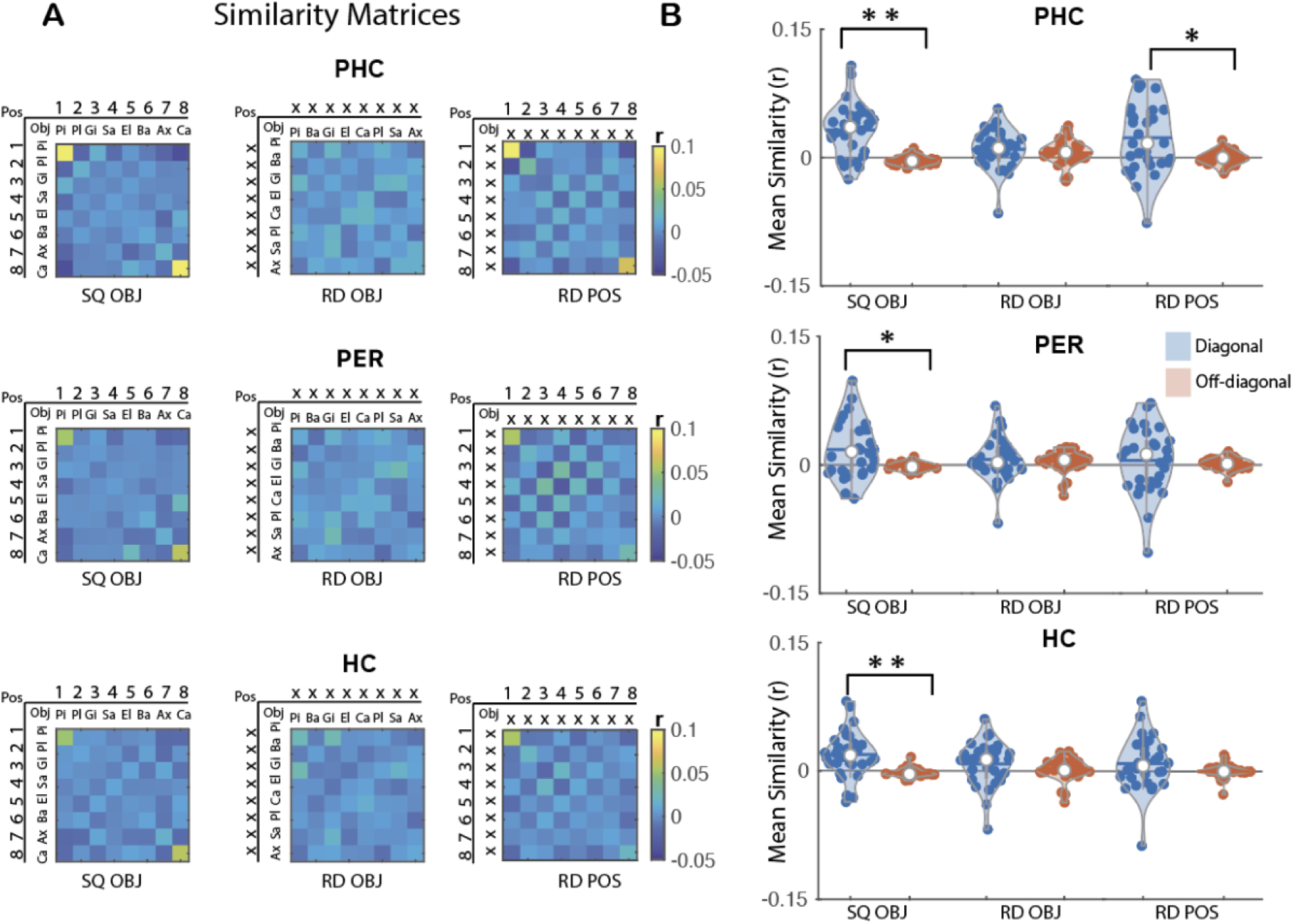
MVPA results for object sequence learning. **A**. Group average neural similarity matrices for the 3 object ROIs. Pattern similarity was computed across repetitions of the object sequence to assess object-position coding in the sequence condition (SQ OBJ matrix, left panel) as well as across repetitions of the random patterns to quantify object coding (RD OBJ matrix, middle panel) and position coding (RD POS matrix, right panel) in the random condition. Color bar represents mean similarity (r). In the POS rows/columns, numbers represent the temporal position in the 8-element sequence or in the random stream. In the OBJ rows/columns, the first 2 letters of each object are presented. (X) represents a random position or object in the object and position matrices, respectively. **B**. Mean pattern similarity for diagonal (blue) and off-diagonal (red) cells as a function of matrices and ROIs. Results indicate that the PHC shows evidence for object-position coding in the sequence condition as well as position coding during random practice. In contrast, PER and HC only show evidence for object-position coding in the sequence condition. Asterisks indicate significant differences between diagonal and off-diagonal (one sided paired sample t-test; Bonferroni corrected \**p_corr_*<.05 and \*\**p_corr_*<.001). Colored circles represent individual data, jittered in arbitrary distances on the x-axis to increase perceptibility. Horizontal lines represent means and white circles represent medians. The shape of the violin [22] depicts the kernel density estimate of the data.

#### Contribution of finger and position coding to finger-position coding

Given that M1 and PMC coded for finger and position/boundary information, it is possible that an overlap in finger and/or position information might contribute to the enhanced pattern similarity for finger-position coding in the learned sequence. To test for this possibility, we performed linear regression analyses to examine whether finger and position coding (as assessed during random practice) in M1 and PMC explained pattern similarity effects observed in the sequence condition. As expected, both finger and position/boundary information contributed to finger-position coding in the motor sequence in M1 (key: B=0.90, t(_1,27_)=6.5, *p_corr_*<0.005; pos: B=1.18, t(_1,27_)=4.83, *p_corr_*<0.005) and PMC (key: B=0.77, t(_1,27_)=4.08, *p_corr_*<0.005; pos: B=1.15, t(_1,27_)=9.02, *p_corr_*<0.005, and see Supplementary Table S2 for results on all other ROIs).

#### Summary

Altogether, these results indicate that hippocampal patterns carry information about fingers in their learned temporal position in the motor sequence condition (i.e., finger-position coding). Although M1 and PMC showed evidence of representing finger-position information in the sequence condition, these regions also exhibited finger and position (PMC) or boundary position (M1) coding in the random condition. Linear regressions indicate that finger-position coding in M1 and PMC during the execution of the learned sequence can, at least in part, be attributed to finger and position/boundary.

### Declarative memory domain

As above, within-declarative memory domain analyses first examined *selective coding* in the ROIs. We measured here the extent to which specific ROIs represent information about (1) objects in their learned temporal position in the sequence (i.e., object-position coding in the sequence condition), (2) objects irrespective of their position (i.e., object coding in the random condition), and (3) temporal positions irrespective of the object (i.e., position coding in the random condition). When the ROIs presented evidence of coding for multiple representations (e.g., both object-position and object coding), we performed linear regression analyses to assess the *contribution of the different representations* to object-position coding. Within-declarative memory domain analyses were performed and corrected on all 5 ROIs but only results corresponding to the 3 ROIs involved in object sequence learning (i.e., the perirhinal cortex PER, the parahippocampus PHC and the hippocampus HC) are presented in the main text (see Supplemental Figures S1 and S2 and Table S3 for results related to the other ROIs).

#### Object-position coding in a sequence

To investigate whether the object sequence ROIs represent information about objects in their learned temporal position in the sequence, we computed the correlation in voxel activation patterns between pairs of individual objects across repetitions of the object sequence (Fig. 4A). The resulting 8×8 matrix (***SQ OBJ***, Fig. 4A) revealed significantly higher mean similarity along the diagonal (i.e., same object + position) as compared to the off-diagonal (i.e., different object + position) in all object ROIs (paired sample t-test: PHC, t(29)=5.62, d=1.03, p_corr_<0.005; PER, t(29)=2.95, d=0.54, p_corr_=0.02; HC, t(29)=3.73, d=0.68, p_corr_<0.005, Fig. 4B). These results indicate that all object ROIs carry information about objects in their learned temporal position in a sequence.

#### Object and position coding in the random condition

Similar to above, control analyses were performed using the random data to test whether the ROIs also represented information about object and position. Results derived from the 8×8 ***RD OBJ*** matrix (Fig. 4A) did not reveal any significant differences in mean similarity along the diagonal (i.e., same object) as compared to the off diagonal (i.e., different object) in any of the object ROIs (PHC, t(29)=1.06, d=0.19, p_corr_=0.74; PER, t(29)=0.54, d=0.1, p_corr_=1; HC, t(29)=1.49, d=0.27, p_corr_=0.37, Fig. 4B). The 8×8 ***RD POS*** matrix (Fig. 4A) revealed significantly higher mean similarity along the diagonal (i.e., same position) as compared to the off-diagonal (i.e., different position) for the PHC (t(29)=2.88, d=0.53, p_corr_=0.02) but not for the PER or the HC (PER, t(29)=0.56, d=0.1, p_corr_=1; HPC, t(29)=1.45, d=0.26, p_corr_=0.4). It is worth noting that position coding in the PHC was no longer significant after removing boundary positions from the position matrix (see Supplementary Table S1 for corresponding statistics). These results suggest that PHC carries information about boundary positions.

#### Contribution of position coding to object-position coding

As the PHC presented evidence of object-position coding and boundary position coding, we performed linear regression analyses to examine whether boundary position coding (as assessed during random practice) might explain pattern similarity effects observed in the sequence condition. As expected, boundary position information contributed to the object-position coding effect in the object sequence in the PHC (B=0.49, t(_1,,28_)=4.28, *p_corr_*<0.005 and see Supplemental Table S3 for results in all other ROIs).

#### Summary

Overall, these results indicate that HC and PER patterns carry information about objects in their learned temporal position in the object sequence condition (i.e., object-position coding). Although PHC showed evidence of object-position coding in the sequence condition, this region also exhibited position (edge) coding in the random condition. Linear regression analyses indicated that object-position coding in the PHC during the execution of the learned sequence can, at least in part, be attributed to position/boundary coding.

### Domain-general effects (across memory domains)

As above, between-memory domains analyses first examined *selective coding* in the ROIs. We measured here the extent to which specific ROIs represent information about (1) item (irrespective of their nature) in their learned temporal position in the sequence (i.e., item-position coding in the sequence condition), (2) items irrespective of their nature and their position (i.e., item coding in the random condition), and (3) temporal positions irrespective of the item (i.e., position coding in the random condition). When the ROIs presented evidence of coding for multiple representations (e.g., both item-position and position coding), we performed linear regression analyses to assess the *contribution of the different representations* to item-position coding across domains.

#### Item-position coding across sequences of different domains

We tested whether the representation of temporal order information during the execution of learned sequences is content-free. To do so, we examined whether items from different domains (i.e., fingers and objects) but sharing the same temporal position in the different sequences (e.g., key X in position 2 in the motor sequence and object Y in position 2 in the object sequence) elicited similar multivoxel brain patterns. We therefore build a representational similarity matrix in which voxel activation patterns were correlated between the object and motor sequence tasks. The resulting 8×8 matrix (***SQ ACROSS***, Fig. 5A) revealed significantly higher mean similarity along the diagonal (i.e., object/key + same position) as compared to the off-diagonal (i.e., object/key + different position) in all ROIs (paired sample t-test: M1, t(29)=6.97, d=1.27, p*_corr_*<0.005; PMC, t(29)=7.96, d=1.45, p*_corr_*<0.005; HC, t(29)=5.61, d=1.02, p*_corr_*<0.005; PHC, t(29)=6.76, d=1.23, p*_corr_*<0.005; PER, t(29)=6.26, d=1.14, p*_corr_*<0.005, Fig. 5B). These results indicate that all ROIs carry information about items (irrespective of their domain) in their learned position in the sequence.

**Figure 5.**
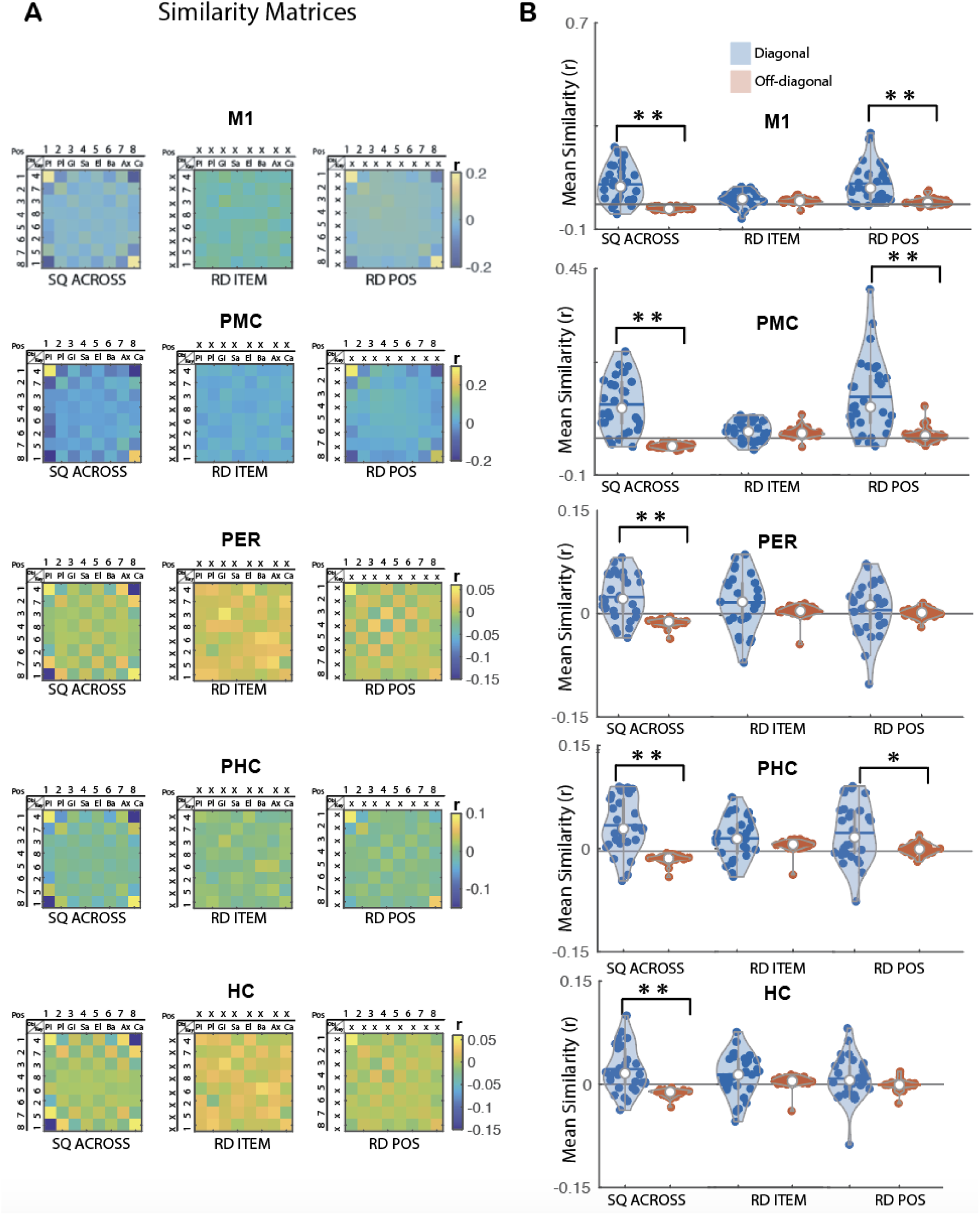
MVPA results for sequence learning across memory domains. **A**. Group average neural similarity matrices for all 5 ROIs. Pattern similarity was computed between pairs of the individual objects from the object sequence condition and individual key presses from the motor sequence condition to assess item-position coding across sequence conditions (SQ ACROSS matrix, left panel) as well as across repetitions of the random patterns to quantify item coding (RD ITEM matrix, middle panel) and position coding (RD POS matrix, right panel) in the random condition. Color bar represents mean similarity (r). In the POS rows/columns, numbers represent the temporal position in a the 8-element sequences or the random stream. In the KEY rows/columns, numbers represent fingers. In the OBJ rows/columns, the first 2 letters of each object are presented. (X) represents a random position or key/object in the item and position matrices, respectively. **B**. Mean pattern similarity for diagonal (blue) and off-diagonal (red) cells as a function of matrices and ROIs. Results indicate that M1, PMC and PER show evidence for item-position coding in the sequence condition and position coding while HC and PER only represent item-position coding across sequences. Asterisks indicate significant differences between diagonal and off-diagonal (one sided paired sample t-test; Bonferroni corrected \**p_corr_*<.05 and \*\**p_corr_*<.001). Colored circles represent individual data, jittered in arbitrary distances on the x-axis to increase perceptibility. Horizontal lines represent means and white circles represent medians. The shape of the violin [22] depicts the kernel density estimate of the data. Note that as in earlier research [16], Y axis scales are different between ROIs to accommodate for differences in signal-to-noise ratio (and therefore in effect sizes) between ROIs.

#### Item and position coding in the random condition

Similar as above, control analyses were performed using the random data to test whether the ROIs also represented information about item across domains (i.e., similarity between e.g., key X and object Y that were presented in the same temporal position in the sequence condition) and position information (as above). Results derived from the ***RD ITEM*** 8×8 matrix (Fig. 5A) did not reveal any significant differences in mean similarity along the diagonal (i.e., object and key in the random condition but that were presented in the same temporal position in sequence condition) as compared to the off diagonal (i.e., objects and keys with different temporal positions in the sequence) in any of the ROIs (M1, t(29)=1.04, d=0.19, p*_corr_*=0.77; PMC, t(29)=0.52, d=0.1, p*_corr_*=1; HC, t(29)=1.64, d=0.3, p*_corr_*=0.28; PHC, t(29)=1.86, d=0.34, p*_corr_*=0.19; PER, t(29)=1.94, d=0.35, p*_corr_*=0.16, Fig. 5B). As a reminder, position coding results described above (8×8 ***RD POS***, Fig. 5A) indicate that PMC carries information about position in random patterns and that both M1 and PHC represent information about boundary position in random patterns.

#### Contribution of position coding to item-position coding across sequences of different domains

Given that M1, PMC and PHC presented evidence of item-position coding across sequences of different domains and also coded for position/boundary, we performed linear regression analyses to examine whether the position/boundary coding (as assessed during random practice) in these ROIs might explain pattern similarity effects observed in the sequence condition across memory domains. As expected, position/boundary information contributed to the item-position coding across memory domains in these ROIs (M1, B=0.89, t(_1,,28_)=7.37, *p_corr_*<0.005; PMC, B=0.72, t(_1,,28_)=9.39, *p_corr_*<0.005; PHC, B=0.51, t(_1,,28_)=3.75, *p_corr_*<0.005, see Supplementary Table S4 for results on all other ROIs).

#### Summary

Overall, these results indicate that HC and PER multivoxel patterns carry information about items - irrespective of their memory domain - in their learned temporal position in the sequence condition (i.e., item-position coding). In other words, the patterns associated to a finger are similar to those associated to an object if these items are presented in the same temporal position in a learned sequence. Although M1, PMC and PHC showed evidence of item-position coding across sequences from different domains, these regions also exhibited position (PMC) or boundary position (M1 and PHC) coding in the random condition. Linear regression analyses indicated that item-position coding in these regions during the execution of the learned sequence can, at least in part, be attributed to position/boundary coding.

## Discussion

In the current study, we used multivoxel representational similarity analyses of fMRI data acquired during the execution of learned sequences from different memory domains to test whether the brain processes supporting temporal order coding of sequential behaviors are domain-specific or -general. Our results indicate that the hippocampus and the perirhinal cortex represent information about items in their learned temporal position within both memory domains separately, but also across memory domains, i.e., regardless of whether the item in a particular position is a finger or an object. These regions did not show evidence of finger/object, item or position coding during random practice. In contrast, in all other ROIs, pattern similarity results in the sequence condition within and between memory domains were, at least in part, explained by finger and/or (boundary) position coding assessed in the random condition. Specifically, multivoxel patterns in the primary motor cortex and the premotor cortex represented both finger- and (boundary) position-based information during random practice while the parahippocampus carried information about boundary positions.

Our results indicate that hippocampal multivoxel patterns carried information about the position of items in a learned sequence within both memory domains but also across sequences from different memory domains. These results suggest that the hippocampus similarly encodes the temporal order of items in a sequence irrespective of their nature. Importantly, and in contrast to other ROIs (but see discussion on the perirhinal cortex below), the hippocampus did not represent information about finger/object, item or position during random practice. The observation of item-position coding in a learned sequence within each memory domain is consistent with earlier multivariate fMRI studies demonstrating that the hippocampus represents information about the temporal position of objects [15] and movements [16] in learned sequences, but not about the item (object/finger) or temporal position in random patterns. It is also consistent with evidence suggesting that the hippocampus processes temporal order information during sequence learning irrespective of the nature of the items (e.g., letters, odors, objects and movements; [15,16,18,23] and the type of memory (i.e., motor vs. non-motor: [16]; [15]; spatial vs. non-spatial: [24–25]. Importantly, we show here that hippocampal multivoxel patterns are similar between items from different domains (i.e., pattern associated to a movement vs. an object) when they are presented in the same ordinal position in a learned sequence. This is in line with earlier views proposing that the hippocampus represents the associative map linking response goals - rather than item-specific information - to their ordinal position in a sequence [16]. The current data suggests that such map, abstracted during sequence learning, is independent of the type of memory items to order and might serve as a cognitive spatiotemporal framework to order sequential behaviors irrespective of their domains. This concurs with the sequence-generator hypothesis, an influential model of hippocampal functioning suggesting that the hippocampus provides a sequential, content-free, structure that preserves the order of experiences irrespective of their nature [19]. Altogether, these results provide the first experimental evidence that the capacity of the hippocampus to encode temporal order during sequence learning is a general process that is shared across memory domains.

Albeit unexpected, a similar pattern of results was observed in the perirhinal cortex. Specifically, perirhinal multivoxel patterns of activity represented information about items in their learned temporal position in an object sequence, in a motor sequence (see Supplemental Figure S2) and regardless of the type of sequence while no item nor position coding was observed. This contradicts earlier research showing that the perirhinal cortex represents information about objects [15,26,27] but not about objects in their learned temporal position in a sequence [15]. The reason for these discrepancies is unclear but one could argue that they are related to the visual stimulation paradigm used in our experiment. Our task necessitated to not only present the central object (visual cue triggering movement) on the screen but also to constantly display the object-key mapping at the bottom of the screen. This resulted in the simultaneous presentation of multiple objects on the screen which may have masked the representation of the central cued objects in the perirhinal cortex. While this is possible, we argue that this is unlikely as we were able to observe the well-known object-category representation in the ventral occipito-temporal cortex (e.g., [28,29], see supplemental Figure S3) which suggests successful visual processing of the central image in this brain area. Based on these results, we cannot completely rule out that object coding did not contribute to item-position coding in the perirhinal cortex but the current pattern of results suggest that, similar to the hippocampus, the perirhinal cortex is specifically involved in associating memory items to their learned position in a sequence regardless of the nature of the memory item. These findings are in contrast with earlier work suggesting that the perirhinal cortex is involved in representing *domain-specific* relational binding information that is fed into the hippocampus for *domain-general* processing [30]. However, they are line with a growing body of literature showing *domain-general* effects in the perirhinal cortex for the encoding of relationships between items and events [31], including item-item [32], item-time [33] and item-location [34] associations. We therefore propose that both the perirhinal cortex and the hippocampus have a similar capacity to represent memory items in their learned temporal position in a sequence, irrespective of the nature of the items to order.

In contrast to the medial temporal lobe regions discussed above, the primary motor cortex, the premotor cortex and the parahippocampus carried information about multiple representations (i.e., item-position and item/position coding). The results of the linear regression analyses indicated that the contribution of these brain regions to sequence learning across domains is attributed - at least in part - to the coding of some of these overlapping representations. In the premotor cortex, position coding significantly contributed to item-position coding across sequences of different domains. The role of the premotor cortex in temporal processing is well described in the motor domain. For example, animal work has shown that premotor cortex neurons are tuned to specific positions in a series of movements [35] and multivariate analyses of human fMRI data indicate that the premotor cortex encodes temporal features about repetitive movements irrespective of their patterns (different sequences, [36]; sequence and random patterns, [16]). The premotor cortex is therefore thought to provide a temporal scaffold in which movements are implemented during motor task practice [16]. We are not aware of any report of such function of the premotor cortex in the declarative memory domain (but note that the premotor cortex is usually not considered a region of interest in this memory domain). We therefore argue that the involvement of the premotor cortex in position coding across domains in the current study (as well as within each memory domain, see supplemental Figure S2 and Tables S2-S4) is related to the motoric nature of the task rather than to a temporal processing function shared across memory domains. Indeed, all task variants (including the object sequence task) required participant to respond to each cue with a key press. In line wither earlier work [16,36], we suggest that the involvement of the premotor cortex is therefore presumably related to the implementation of a temporal scaffold in which movements are implemented irrespective of the version (motor vs. object) and sequential nature (sequence vs. random, respectively) of the task. This, however, remains speculative.

In the primary motor cortex, boundary position coding significantly contributed to item-position coding across domains. The boundary effects observed in this study are consistent with our earlier research showing greater similarity values at the start and end of the stream of 8 elements (in both motor sequence and random conditions) across all multiple regions of interest including the primary motor cortex [16]. It is worth noting that these boundary effects are particularly pronounced in our study in comparison to our earlier work [16]. We hypothesize that this is related to the change in object/key mapping that occurred after each stream of 8-elements and that was designed to orthogonalize motor and object sequence tasks in the current study. This might have accentuated attentional [37] and anticipatory [38] processes in the primary motor cortex; processes that are known to increase the similarity of brain responses [39]. We therefore suggest that the contribution of the primary motor cortex to item-position coding across memory domains is related to non-memory related attentional processes that were particularly pronounced in this paradigm.

Similar to the primary motor cortex, boundary position coding in the parahippocampus significantly contributed to item-position coding across memory domains. Earlier research has shown that the parahippocampus codes for the temporal position of objects presented in a stream irrespective of whether they are presented in a sequential or random order [15]. It is also suggested that the parahippocampus carries information about the context in which objects are encountered [40,41]. Previous MEG research also indicated such temporal function in the parahippocampus during motor learning and showed that this region is involved in coding an abstract template of ordinal position during motor sequence preparation [42]. Together with the evidence reviewed above demonstrating parahippocampal involvement in temporal coding in both the declarative and the procedural memory domains, the current findings suggest that the role of the parahippocampus in position coding might be shared across memory domains. However, it is important to note that, similar to the primary motor cortex, position coding in our dataset was no longer observed in the parahippocampus after removing boundary positions. It is therefore possible that similar attentional processes as discussed above might have contributed to the pattern of results in the parahippocampus. Altogether, while our data and earlier research both point to similar position coding effect in the parahippocampus between memory domains, it cannot be ruled out that these effects are related to boundary positions only.

Last, the current within-motor memory domain results partly confirm our earlier research [16]. Specifically, our data indicate that the contribution of motor cortical regions to motor sequence learning is attributed - at least in part - to movement and position coding. Specifically, in the primary motor cortex, both finger and boundary position information contributed to finger-position coding. The contribution of finger information to finger-position coding is consistent with earlier multivariate fMRI studies demonstrating that although the primary motor cortex receives input from areas showing motor sequence representation, multivoxel activity patterns in this region relate to the execution of individual key presses during motor sequence learning [16,43]. The contribution of boundary position coding to finger-position coding in the primary motor cortex was not observed in our earlier research [16] and might be attributed to the nature of the current paradigm (see above). In the premotor cortex, both finger and position (beyond boundaries) information contributed to finger-position coding. The observation of finger coding in the premotor cortex contradicts earlier research indicating that this region encodes sequential context rather than individual movement components [43–45]. However, they are in line with our recent work [16] suggesting that the premotor cortex represents finger information during early stages of learning (as in the current study) while sequence representations may emerge at later learning stages [43,45]. As discussed above, our results point to a role of the premotor cortex in temporal position coding which is in line with both previous animal [35] and human [16,36] research and speak to the capacity of the premotor cortex to provide a temporal scaffold in which movements are implemented.

A limitation in our study is that, except for the MRI session, participants performed significantly better in the motor sequence task than in the object sequence task. We speculate that these differences are related to the nature of the different tasks and do therefore not necessarily reflect difference in learning amplitude between tasks. Specifically, as learning progressed in the motor version of the task, participants might have been able to press the sequence of keys without relying on the key-object mapping displayed on the screen. In contrast, the object sequence task required participants to use the key/object mapping to respond even at later stages of learning. This hypothesis is supported by the data from the MRI sessions that were acquired with longer response-stimulus-interval (see methods) and during which similar performance was observed between motor and object sequence conditions, presumably due to the additional time available for the mapping that was particularly beneficial for the object task. As earlier research suggests that performance speed can alter brain representations [44], we performed additional control analyses to assess potential relationships between performance and brain patterns. Specifically, we tested whether the level of performance – and potentially the strength of the memory trace – reached at the end of training on day 1 was related to the strength of the motor and object sequence representations assessed during the MRI session the next day (see section 3 of the supplementary results). Our results showed no correlation between performance and finger/object-position coding in any of the ROIs, suggesting that the difference in performance observed between tasks on day 1 is unlikely to influence the pattern of brain results reported above.

To conclude, our findings suggest that medial temporal lobe regions such as the hippocampus and the perirhinal cortex play a critical role in representing memory items - irrespective of their domains - in their learned temporal positions in a sequence. We propose that these regions contribute to the development of domain-general item-position maps that provide a content-free cognitive framework to order sequential behaviors irrespective of their nature. In line with the sequence-generator hypothesis [19], our results support the view that the medial temporal lobe provides a series of indices that point to cortical modules carrying content-specific information (e.g., primary motor cortex for finger information and premotor cortex for position information). Our findings concur with the hypothesis that these different task-specific inputs are sequenced by activity patterns in the medial temporal lobe, thus preserving the ordinal structure over which experiences occur, irrespective of their nature.

## Material and Methods

### Participants

Sample size computation was done with G*Power [46]. The effect size was calculated based on previous work on motor memory [16] that detected significant differences in hippocampal activation patterns between conditions within the motor domain (i.e., 3 conditions: finger-position, finger and position information; main effect of condition; effect size = 0.40). The correlation coefficient was set to 0.5 for a more conservative power calculation to account for the different nature of the present design (2 tasks; 6 conditions). Sphericity correction was set to 0.5 to account for potential deviation from the sphericity assumption. Alpha was set to 0.01 to correct for the number of ROIs (5) and a power of 0.8. The required sample size was 28 participants.

To account for attrition, 33 young healthy adults aged 20-35 (18 female, mean age=26.3, SD=5.2) were recruited to participate in this study. All participants completed an online screening questionnaire to verify their eligibility to take part in the study. All participants were right-handed (Edinburgh Handedness Questionnaire; [47], self-reported no current or previous history of neurological, psychological, or psychiatric conditions, and were free of any psychotropic or sleep impacting medications. None of the participants received previous extensive training as a musician or as a professional typist. They also presented no contra-indication to MRI. Further screening using standardized questionnaires also ensured that participants did not exhibit any indication of anxiety (Beck Anxiety Inventory (BAI); [48]) or depression (Beck Depression Inventory (BDI); [49]). Participants reported normal sleep quality and quantity during the month prior to the study (Pittsburgh Sleep Quality Index (PSQI); [50]) as well as normal levels of daytime sleepiness (Epworth Sleepiness Scale; [51]). None of the participants were extreme evening or morning type individuals (Morningness-Eveningness Questionnaire (MEQ); [52]). A total of 30 participants were included in the analysis. One participant was excluded for low performance accuracy on the sequence tasks (below the mean – 3 standard deviations from the group), another for excessive movement in the MRI scanner across multiple runs (> 2 voxels, see below for movement information at the group level) and another for a non-compliant sleep schedule between experimental sessions (slept less than 7 hours and went to sleep later than 1am). All study procedures were approved by the University of Utah institutional review board (IRB_00155080). All participants provided written informed consent at the start of the study and received monetary compensation for their time. Participants’ demographics as well as information on their sleep and vigilance are presented in supplementary Table S5.

### Serial Reaction Time Task (SRTT)

All participants performed an adapted version of the explicit Serial Reaction Time Task (SRTT; [3] which was coded in MATLAB (Mathworks Inc., Sherbom, MA) using Psychophysics Toolbox version 3 [53]. The task consisted of an 8-choice reaction time task in which participants were instructed to react to visual cues shown on a screen by pressing a corresponding key on a keyboard (behavioral sessions) or on an MRI-compatible button box (MRI session). The large visual cue presented in the middle of the screen corresponded to one of eight different images of objects (Fig. 1A, left panel). The 8 different objects were constantly presented on the bottom of the screen during task practice and mapped to each of the eight fingers used to perform the task (i.e., no thumbs). Participants were instructed to respond to the central visual cue as fast and as accurately as possible by pressing the corresponding key according to the object/key mapping displayed at the bottom of the screen (see Fig. 1A, left panel). When the response was recorded, the central cue turned to a green fixation cross while the mapping with the 8 objects remained on the bottom of the screen during the full duration of the response-stimulus-interval (RSI). RSI was 750ms for the behavioral version of the task and jittered between 1.5-2.5s for the MR version of the task in order to optimize the design for MVPA (see procedure and MVPA sections below). Blocks of task practice were alternated with 10s rest periods which were indicated by a red fixation cross in the center of the screen (replacing the central cue object / green cross displayed during practice) and red frames replaced the locations of the 8 objects that displayed the key-object mapping on the bottom of the screen. Participants were made aware that the order of objects/movements would follow a repeating sequential pattern but were not told the precise sequence or the number of elements.

Importantly, the key-object mapping displayed on the bottom of the screen changed after each stream of 8 elements (i.e., every 8 objects / 8 key presses) in order to orthogonalize motor and object sequence learning tasks. With this manipulation, we were able to develop three different versions of the task. In the *object sequence condition*, the stream of objects on the screen followed a sequential pattern (object sequence, e.g., pineapple, plane, giraffe, saw, elephant, banana, axe, car) while the responses on the keyboard followed a random order (motor random). In the *motor sequence condition*, the responses on the keyboard followed a repetitive sequential pattern (motor sequence, e.g., 4,7,3,8,6,2,5,1, whereby 1 and 8 represent the left and right little fingers, respectively) while the stream of objects on the screen followed a random order (object random). In the *random task condition*, the series of both responses and objects followed a random order (i.e., motor random / object random). This experimental manipulation allowed us to examine sequence learning in the two different memory domains using the same task. Note that all participants learned the same object and motor sequences. However, to optimize the MVPA of the fMRI data, the starting point of the sequence in each task condition was different across participants such that the same key/object was not associated to the same temporal position in a sequence across individuals (i.e., in participant 1, the sequences started with 4/pineapple, in participant 2, the sequences started with 7/plane, etc.). The random task variant served 3 functions. The first was to ensure that general motor execution, independent of sequence learning *per se* (e.g., baseline performance levels) did not differ between the different experimental sessions. The second was to assess sequence-specific learning on Day 2 (i.e., comparison between sequence and random conditions). The third was to serve as a neuroimaging control condition that allowed us to investigate which brain areas coded for temporal position (irrespective of the memory type) but also for specific task items (i.e., finger- and object-coding, see MVPA section below). In all task variants, the number and timing of key presses was recorded and used to compute behavioral outcomes (see below).

### Procedure

The experimental design is presented in Fig. 1B. Participants were asked to complete 3 experimental sessions spread over 2 consecutive days. They were instructed to follow a constant sleep schedule (according to their own schedule ±1h; latest bedtime: 1am; minimum 7h of sleep; no naps) starting 3 days before the first session and between the two experimental days. Compliance to the sleep schedule was monitored using sleep diaries and wrist actigraphy (ActiGraph wGT3X-BT, Pensacola, FL). On day 1, participants learned the motor and object sequence tasks during 2 separate 1hr sessions (order counterbalanced across participants) performed outside the MRI scanner. The first session was completed in the morning (between 9am-12pm) while the second session was performed at least 4 hours later (i.e., between 1-4pm) in order to reduce interference between the 2 learning episodes [54]). During session 1, participants completed a brief questionnaire to collect information regarding their sleep patterns in the previous 24 hours (St Mary’s Hospital Sleep Questionnaire (SMS); [55]) their subjective feelings of alertness (Stanford Sleepiness Scale (SSS); [56]) and any consumption of alcohol and/or recreational drugs in the last 24 hours. Next, participants performed a psychomotor vigilance test (PVT) to obtain an objective measure of their alertness. For the PVT, participants were asked to fixate on a cross in the center of the screen and to press the space bar as fast as possible when the fixation cross disappeared. A timer appeared on the screen displaying the participants’ reaction time in milliseconds. Participants completed 50 trials of the PVT task. Next, participants were instructed to practice the SRT task with an RSI of 750ms (behavioral version of the task). Practice was organized in blocks of 5 repetitions of an 8-element series. First, participants completed 4 blocks of the random condition to get familiar with the task apparatus and measure baseline performance. They were then trained on 20 blocks of either the object or motor sequence condition. After a short break offered to minimize the confounding effect of fatigue on end-training performance [57], performance was tested with 4 additional blocks of practice.

After completing session 1, participants were given actigraph monitors to be worn until they returned for the final session on day 2. For the 4 hours between sessions 1 and 2, participants were instructed to not take a nap or practice the task and to avoid intense exercise and consumption of stimulating substances (e.g., alcohol, caffeine, energy drinks). Session 2 followed the same procedure as session 1 except that participants were trained on the other sequential condition (object or motor). After completing session 2, participants were instructed to have a regular night of sleep according to the schedule followed prior to the experiment, to continue wearing the actigraph, to not practice the task and avoid consumption of stimulating substances. Participants returned 24 hours later, at the MRI facility, for the third and final session.

On experimental day 2, for session 3, participants were asked to complete the same questionnaires that were completed on day 1 along with an MRI screening form. Next, they performed the PVT followed by 1 practice block of familiarization to the MR version of the SRTT task optimized for MVPA (slower RSI; mixed conditions; different rest pattern; see below) outside the scanner. Participants were then placed in the MR scanner so that their brain activity was recorded while performing the task on an MRI-compatible keyboard (Psychology Software Tools, Celeritas). The MRI session consisted of 8 runs (Fig. 1B). Each run lasted approximately 6min and included the 3 following conditions of interest for MVPA: motor sequence, object sequence and random. Each run comprised 5 repetitions of each condition (8 runs x 5 repetitions: 40 repetitions per condition in total) and 3 rest blocks of 10s (to minimize fatigue) that were randomly positioned between conditions. The order of the conditions was pseudo-randomized within each run so that the same condition was not repeated more than two times in a row. The inter-stimulus-interval was jittered between 1500-2500ms. There was no additional delay between conditions. After completing the MR session, participants were tested with 4 additional blocks of each of the 2 sequential conditions followed by a final 4 blocks of the random condition outside the scanner with the behavioral version of the task (see above) to assess motor memory retention.

### fMRI data acquisition

Images were acquired with a Siemens Vida 3.0T MRI System and a 64-channel head coil. Functional images were acquired with a T2* gradient echo-planar sequence using axial slices angled to maximize whole brain coverage (TR = 1.03s, TE = 31 ms, FA = 65°, 56 slices, 2.5 mm isotropic voxels, 0 mm slice gap, FoV = 210 × 210 mm^2^, matrix size = 88 × 88 × 56 slices, MB = 4). A structural T1-weighted 3D MP-RAGE sequence (TR = 2.5s, TE = 2.98 ms, FA = 8°, 176 slices, FoV = 250 × 250 mm^2^, matrix size = 256 × 256 × 176) was also obtained for each participant during the same scanning session.

### Data analysis

#### Behavioral data

Two measures of performance were extracted from the behavioral data. The measure of performance speed was response time (in seconds) calculated as the time between the onset of display of the central cue and the response of the participant on the keyboard. Response time on correct trials were averaged within each practice block. The measure of performance accuracy consisted of percentage of correct responses per block. To assess whether baseline performance differed between sessions on day 1, two separate repeated-measures ANOVA were conducted on performance speed and accuracy on the random SRTT data using session condition (2: object session, motor session) and block (4) as the within-subject factors (see Fig. 2 and corresponding results are reported in the supplemental information). To assess learning over practice blocks and compare learning across conditions on day 1, separate two-way repeated-measures ANOVAs were performed on both speed and accuracy measures using condition (2: object, motor) and block (20 for training, 4 for test) as the within-subject factors. Behavioral data collected on day 2 were analyzed with separate repeated-measures ANOVA using condition (3: object, motor, random) and block (8 for the MRI session and 4 for the retest session) as the within-subject factors. Last, additional control analyses were performed to assess the effect of order on performance during learning on day 1 and retest on day 2. To do so, the analyses described above were repeated using session order, irrespective of the task condition (2: Session 1, Session 2) and block as the within-subject factors. The results corresponding to these analyses are reported in section 2 of the supplemental results. Statistical tests yielding p values ˂ 0.05 were considered significant.

#### fMRI data

##### Preprocessing

Anatomical and functional images were preprocessed using SPM12 (Welcome Department of Imaging Neuroscience, London, UK) implemented in MATLAB. Preprocessing included segmentation of the anatomical image into gray matter, white matter, cerebrospinal fluid (CSF), bone, soft tissue and background. Functional images were slice time corrected (reference: middle slice) and realigned using rigid body transformations, iteratively optimized to minimize the residual sum of squares between each functional image and the first image of each session separately in a first step and with the across-run mean functional image in a second step (mean translation in the three dimensions across the 8 runs: 0.27 mm (+/- 0.11 mm), mean maximum translation in the three dimensions across the 8 runs: 0.91 mm (+/- 0.45 mm)). Movement was considered as excessive when translations exceeded more than 2 voxels in either of the three dimensions for any of the 8 runs. One individual was excluded from data analyses as such excessive movements were observed in more than half of the runs. In two other individuals, excessive movement was observed in one specific run and data could be included after truncating volumes in these runs. Specifically, in one individual, the last 111 functional images (out of 382) of one run were excluded and, in another run, the first 13 out of 387 functional images were excluded. The realigned functional images were then co-registered to the structural T1-image using rigid body transformation optimized to maximize the normalized mutual information between the two images. All analyses were performed in native space to optimally accommodate interindividual variability in the topography of memory representations [28].

##### Regions of interest

The goal of the fMRI analyses was to examine brain patterns elicited in specific regions of interest (ROIs) by sequence and random task practice. The five following (bilateral) ROIs were selected *a priori* based on previous literature describing their critical involvement in motor and object sequence learning [15–16]: the primary motor cortex (M1) and the perirhinal cortex (PER) for item coding within the motor and declarative memory domains, i.e. for finger and object coding, respectively; the premotor cortex (PMC) and the parahippocampus (PHC) for position coding described in motor and object sequence learning, respectively; and the hippocampus (HC) for item-position coding in both domains. As in previous studies, all ROIs were anatomically defined (e.g., [16,58]. The bilateral hippocampal ROI was defined in native space using FSL’s automated subcortical segmentation protocol (FIRST; FMRIB Software Library, Oxford Centre for fMRI of the Brain, Oxford, UK). Bilateral cortical ROIs were created in MNI space using the Brainnetome atlas [59] with probability maps thresholded at 35%. The M1 ROI included areas A4ul (upper limb) and A4hf (hand function) while the PMC comprised areas A6cdl (dorsal PMC) and A6cvl (ventral PMC). The PER included A35_36c (caudal area) and A35_36r (rostral area) as in [60] and PHC comprised both TH (medial posterior PHC) and TL (lateral posterior PHC) areas [59]. The resulting ROI masks were mapped back to individual native space using the individual’s inverse deformation field output from the segmentation of the anatomical image. All ROIs were registered to the functional volumes and masked to exclude voxels outside of the brain. Voxels showing less than 10% grey matter probability were also excluded from the ROIs. The average number of voxels within each ROI is reported in Supplementary Table S6.

##### Representational similarity analyses (RSA)

To assess coding of temporal order and item information during sequence learning within and across memory domains, we used multivoxel pattern analyses of task-related fMRI data. In particular, we used representational similarity analyses (RSA) which are based on the assumption that if a particular type of information is processed by a particular brain region, then it will be represented in the pattern of activation across voxels in that brain region [21]. Such analyses therefore predict higher correlations in voxel activity between pairs of trials sharing this information as compared to pairs of trials not sharing this information. Using RSA on the sequence condition data, we first assessed the extent to which our ROIs represent information about memory items in their learned temporal position in a sequence (1) **within the motor memory domain**, i.e., representation of fingers in their learned position in the motor sequence referred to as *finger-position coding*, (2) **within the declarative memory domain**, i.e. representation of objects in their learned position in the object sequence referred to as *object-position coding*, and (3) **across memory domains**, i.e., representation of items (irrespective of the memory domain) in their learned position in the sequence referred to as *item-position coding*. Next, using the random condition data, we performed control analyses to assess to what extent the different ROIs also represented information about finger/object and position. Specifically, we tested whether the ROIs carry information about (1) fingers irrespective of their temporal position in the stream of movement (i.e., finger coding), (2) objects irrespective of their temporal position in the stream of objects (object coding); and (3) temporal positions irrespective of finger/object information (i.e., position coding).

To perform the RSA, 3 separate GLMs were fitted to the 8 runs of preprocessed functional data of each participant. The first GLM (i.e., position GLM) was constructed to model neural responses evoked for each temporal position and include one regressor for each position (8 temporal positions in the stream) for each of the 3 conditions (motor sequence, object sequence and random) in each run (8 positions x 3 conditions = 24 regressors per run, representing positions in motor sequence, positions in object sequence and positions in random condition; 8 runs, i.e., 192 regressors in total). Similarly, the second GLM (i.e., key GLM) was constructed to model neural responses evoked by individual key presses and include one regressor for each finger/key for each condition in each run (8×3 regressors = 24 regressors per run, representing keys in motor sequence, keys in object sequence and keys in random condition; 8 runs, i.e., 192 regressors in total). Finally, a third GLM (i.e., object GLM) was constructed to model neural responses evoked by individual objects and include one regressor for each object for each condition in each run (8×3 regressors = 24 regressors per run, representing objects in motor sequence, objects in object sequence and objects in random condition; 8 runs, i.e., 192 regressors in total). Since the finger-position and object-position associations are fixed in the object and motor sequence conditions, the same events were modelled in the 3 different GLMs for the sequence conditions. As there is no such association in the random condition, events were reclassified according to (1) temporal position (irrespective of key or object; position GLM), (ii) key pressed (irrespective of temporal position and object; key GLM) or (iii) object (irrespective of key and temporal position; object GLM). For all GLMs, neural responses to events of interest were modelled with delta functions (0ms duration) time locked to cue onsets. Movement parameters and responses during rest periods were entered as nuisance regressors. High-pass filtering with a cutoff of 128 s was used to remove low-frequency drifts from the time series. An autoregressive (order 1) plus white noise model and a restricted maximum likelihood (ReML) algorithm was used to estimate serial correlations in the fMRI signal. The GLMs resulted in separate maps of t-values for each regressor in each run for each ROI. For each voxel within each ROI, the t-values were normalized by subtracting the mean t-value across all regressors within each run separately.

To estimate neural similarity matrices for each ROI, the full dataset (8 runs) was randomly divided into two independent subsets of 4 runs and t-values were averaged within each set to give 2 sets of t-values for each regressor for each voxel. For a given condition within one of the GLMs, the t-values for all the voxels within an ROI from one set was correlated with the second set for all combinations of the 8 regressors for that condition (e.g., position 1-8). This resulted in an 8 x 8 matrix of Pearson correlation coefficients. This procedure was repeated 140 times (i.e., 2 conditions x 70 possible combinations when choosing 4 numbers out of 8), each time averaging over different subsets of runs. The correlation coefficients were fisher-transformed and then averaged across the 140 iterations resulting in an 8 x 8 correlation matrix for each participant, ROI, and research question of interest. A total of 7 matrices were constructed to address the different research questions, i.e., which ROIs code for (1) finger-position coding (within motor sequence condition), (2) object-position coding (within object sequence condition), (3) item-position coding (between motor and object sequence conditions), (4) temporal position (within random condition), (5) key (within random condition) and (6) object (within random condition). Note that an additional control matrix was created to test for (7) item coding (within random condition) to control for the potential contribution of item similarity between memory domains (e.g., any unexpected similarity - during random practice - between e.g., key 2 and elephant items that are presented in the same temporal position in the sequence condition) to the results observed in the item-position matrix (see Figs. 3-5).

We first examined ***selective coding*** in our different ROIs, i.e. whether voxel activity patterns were more similar during the repetition of trials sharing the same information as compared to pairs of trials not sharing this information. Since the diagonal of the correlational matrix represents a correlation between the same information (e.g., key 4 vs. key 4), if a brain region codes for this type of information, then we would expect a high similarity index when compared to correlations between information that is different (off-diagonal, e.g., key 4 vs. key 1). To investigate whether a given brain region coded for a particular type of information, for each ROI and each type of matrix, a one-tailed paired sample t-test was therefore performed across participants to compare diagonal vs. non-diagonal mean similarity.

Next, when the ROIs presented evidence of coding for multiple representations (e.g., item-position in the sequence condition and item coding in the random condition), we assessed the contribution of the different overlapping representations derived from random practice to the coding results observed in the sequence condition using forward stepwise linear regression analyses. Specifically, the goal of these analyses was to identify whether item (finger, object, item) and/or position coding (as assessed during random practice) can explain pattern similarity effects in the sequence conditions. To do so, we ran separate models using delta similarity (i.e., mean pattern similarity diagonal minus off-diagonal) for finger-position/object-position/item-position coding as a dependent variable and delta similarity for finger/object/item and position matrices as predictors. For each ROI, predictors were included in the model only if the ROI showed significant coding for the corresponding information (i.e., a significant difference between diagonal vs. off-diagonal). When two predictors were included in the model, an F-test was used to assess whether the change in explained variance (i.e., R^2^ change; Δ R^2^) from the prior step was significant. Full models including the two predictors within each memory domain and across memory domains can be found for each ROI in supplemental Tables S2-S4.

For all the statistical analyses described above, we applied Bonferroni correction to correct for the 5 ROIs. Note that for motor and object sequence learning, the results reported in the main text only included the 3 memory-domain specific ROIs (i.e., 3 motor ROIs: M1, PMC and HC for motor sequence and 3 object ROIs: PER, PHC and HC for object sequence) but still corrected for 5 ROIs. The supplemental information (Fig. S1-S2) provides results for all ROIs within each memory domain (e.g., motor sequence learning RSA on object ROIs) for completeness.

##### Supplementary analyses

As performance differed between task conditions during initial training (see results section), we examined whether the level of performance reached at the end of the training session on day 1 was correlated with neural pattern similarity examined the next day (note though that performance did not differ between conditions during the scanning session). These control analyses are presented in the Supplementary results section 3 and indicate that the level of performance reached on day 1 is unlikely to influence the pattern of brain results observed the next day.

Additionally, as we did not observe object coding in any of our object ROIs, we performed supplemental analyses to test whether there was successful visual processing of the central object image. Specifically, we tested whether the well-known object-category representation described in the ventral occipito-temporal cortex VOTC (e.g., [28–29]) could be observed in our dataset. To do so, we computed the object representational similarity matrix from the random data (8×8 ***RD OBJ*** matrix) in a VOTC ROI that was defined using the Brainnetome atlas [59] by combining fusiform (37mv, 37lv, A20rv) and infero-temporal areas (A20iv, A37elv, A20r, A20il, A37vl, A20cl, A20cv). As for the other object ROIs, object coding was assessed with the comparison between the mean similarity averaged along the diagonal (i.e., same object) and mean similarity averaged from all off-diagonal cells (i.e., different object). We also assessed object-category coding and compared mean similarities between objects belonging to the same categories (e.g., similarity between the two fruits; pineapple and banana) with the mean similarity between objects belonging to different categories (e.g., similarity between banana and plane). We also compared similarity values averaged across different object categories to similarity within each object category. These results are presented in supplemental Figure S3.

## References

[1] Cohen NJ, Squire LR. Preserved learning and retention of pattern-analyzing skill in amnesia: dissociation of knowing how and knowing that. Science 1980;210:207–10. 10.1126/science.7414331.

[2] Reber PJ, Squire LR. Parallel brain systems for learning with and without awareness. Learn Mem 1994;1:217–29. 10.1101/lm.1.4.217.

[3] Nissen MJ, Bullemer P. Attentional requirements of learning: Evidence from performance measures. Cogn Psychol 1987;19:1–32. 10.1016/0010-0285(87)90002-8.

[4] Albouy G, Fogel S, Pottiez H, Nguyen VA, Ray L, Lungu O, et al. Daytime sleep enhances consolidation of the spatial but not motoric representation of motor sequence memory. PLoS One 2013;8:e52805. 10.1371/journal.pone.0052805.

[5] Henke K. A model for memory systems based on processing modes rather than consciousness. Nat Rev Neurosci 2010;11:523–32. 10.1038/nrn2850.

[6] Schendan HE, Searl MM, Melrose RJ, Stern CE. An fMRI study of the role of the medial temporal lobe in implicit and explicit sequence learning. Neuron 2003;37:1013–25. 10.1016/s0896-6273(03)00123-5.

[7] Albouy G, Sterpenich V, Balteau E, Vandewalle G, Desseilles M, Dang-Vu T, et al. Both the hippocampus and striatum are involved in consolidation of motor sequence memory. Neuron 2008;58:261–72. 10.1016/j.neuron.2008.02.008.

[8] Gheysen F, Van Opstal F, Roggeman C, Van Waelvelde H, Fias W. Hippocampal contribution to early and later stages of implicit motor sequence learning. Exp Brain Res 2010;202:795–807. 10.1007/s00221-010-2186-6.

[9] Albouy G, Fogel S, King BR, Laventure S, Benali H, Karni A, et al. Maintaining vs. enhancing motor sequence memories: respective roles of striatal and hippocampal systems. Neuroimage 2015;108:423–34. 10.1016/j.neuroimage.2014.12.049.

[10] Jacobacci F, Armony JL, Yeffal A, Lerner G, Amaro E Jr, Jovicich J, et al. Rapid hippocampal plasticity supports motor sequence learning. Proc Natl Acad Sci U S A 2020;117:23898–903. 10.1073/pnas.2009576117.

[11] Buch ER, Claudino L, Quentin R, Bönstrup M, Cohen LG. Consolidation of human skill linked to waking hippocampo-neocortical replay. Cell Rep 2021;35:109193. 10.1016/j.celrep.2021.109193.

[12] King BR, Gann MA, Mantini D, Doyon J, Albouy G. Persistence of hippocampal and striatal multivoxel patterns during awake rest after motor sequence learning. IScience 2022;25:105498. 10.1016/j.isci.2022.105498.

[13] Schapiro AC, Reid AG, Morgan A, Manoach DS, Verfaellie M, Stickgold R. The hippocampus is necessary for the consolidation of a task that does not require the hippocampus for initial learning. Hippocampus 2019;29:1091–100. 10.1002/hipo.23101.

[14] Mylonas D, Schapiro AC, Verfaellie M, Baxter B, Vangel M, Stickgold R, et al. Maintenance of procedural motor memory across brief rest periods requires the hippocampus. J Neurosci 2024;44:e1839232024. 10.1523/JNEUROSCI.1839-23.2024.

[15] Hsieh L-T, Gruber MJ, Jenkins LJ, Ranganath C. Hippocampal activity patterns carry information about objects in temporal context. Neuron 2014;81:1165–78. 10.1016/j.neuron.2014.01.015.

[16] Dolfen N, Reverberi S, Op de Beeck H, King BR, Albouy G. The hippocampus represents information about movements in their temporal position in a learned motor sequence. J Neurosci 2024;44:e0584242024. 10.1523/JNEUROSCI.0584-24.2024.

[17] Ezzyat Y, Davachi L. Similarity breeds proximity: pattern similarity within and across contexts is related to later mnemonic judgments of temporal proximity. Neuron 2014;81:1179–89. 10.1016/j.neuron.2014.01.042.

[18] Kalm K, Davis MH, Norris D. Individual sequence representations in the medial temporal lobe. J Cogn Neurosci 2013;25:1111–21. 10.1162/jocn_a_00378.

[19] Buzsáki G, Tingley D. Space and time: The hippocampus as a sequence generator. Trends Cogn Sci 2018;22:853–69. 10.1016/j.tics.2018.07.006.

[20] Friston K, Buzsáki G. The functional anatomy of time: What and when in the brain. Trends Cogn Sci 2016;20:500–11. 10.1016/j.tics.2016.05.001.

[21] Kriegeskorte N, Mur M, Bandettini P. Representational similarity analysis - connecting the branches of systems neuroscience. Front Syst Neurosci 2008;2:4. 10.3389/neuro.06.004.2008.

[22] Bechtold B. Violin plots for Matlab. Github Project 2016;10.

[23] Fortin NJ, Agster KL, Eichenbaum HB. Critical role of the hippocampus in memory for sequences of events. Nat Neurosci 2002;5:458–62. 10.1038/nn834.

[24] Shahbaba B, Li L, Agostinelli F, Saraf M, Cooper KW, Haghverdian D, et al. Hippocampal ensembles represent sequential relationships among an extended sequence of nonspatial events. Nat Commun 2022;13:787. 10.1038/s41467-022-28057-6.

[25] Davidson TJ, Kloosterman F, Wilson MA. Hippocampal replay of extended experience. Neuron 2009;63:497–507. 10.1016/j.neuron.2009.07.027.

[26] Murray EA, Richmond BJ. Role of perirhinal cortex in object perception, memory, and associations. Curr Opin Neurobiol 2001;11:188–93. 10.1016/s0959-4388(00)00195-1.

[27] Clarke A, Tyler LK. Object-specific semantic coding in human perirhinal cortex. J Neurosci 2014;34:4766–75. 10.1523/JNEUROSCI.2828-13.2014.

[28] Haxby JV, Gobbini MI, Furey ML, Ishai A, Schouten JL, Pietrini P. Distributed and overlapping representations of faces and objects in ventral temporal cortex. Science 2001;293:2425–30. 10.1126/science.1063736.

[29] Mattioni S, Rezk M, Battal C, Bottini R, Cuculiza Mendoza KE, Oosterhof NN, et al. Categorical representation from sound and sight in the ventral occipito-temporal cortex of sighted and blind. Elife 2020;9. 10.7554/eLife.50732.

[30] Davachi L. Item, context and relational episodic encoding in humans. Curr Opin Neurobiol 2006;16:693–700. 10.1016/j.conb.2006.10.012.

[31] Suzuki WA, Naya Y. The perirhinal cortex. Annu Rev Neurosci 2014;37:39–53. 10.1146/annurev-neuro-071013-014207.

[32] Messinger A, Squire LR, Zola SM, Albright TD. Neuronal representations of stimulus associations develop in the temporal lobe during learning. Proc Natl Acad Sci U S A 2001;98:12239–44. 10.1073/pnas.211431098.

[33] Naya Y, Suzuki WA. Integrating what and when across the primate medial temporal lobe. Science 2011;333:773–6. 10.1126/science.1206773.

[34] Yanike M, Wirth S, Smith AC, Brown EN, Suzuki WA. Comparison of associative learning-related signals in the macaque perirhinal cortex and hippocampus. Cereb Cortex 2009;19:1064–78. 10.1093/cercor/bhn156.

[35] Merchant H, Pérez O, Zarco W, Gámez J. Interval tuning in the primate medial premotor cortex as a general timing mechanism. J Neurosci 2013;33:9082–96. 10.1523/JNEUROSCI.5513-12.2013.

[36] Kornysheva K, Diedrichsen J. Human premotor areas parse sequences into their spatial and temporal features. Elife 2014;3:e03043. 10.7554/eLife.03043.

[37] Bhattacharjee S, Kashyap R, Abualait T, Annabel Chen S-H, Yoo W-K, Bashir S. The role of primary motor cortex: More than movement execution. J Mot Behav 2021;53:258–74. 10.1080/00222895.2020.1738992.

[38] Lee CS, Aly M, Baldassano C. Anticipation of temporally structured events in the brain. Elife 2021;10. 10.7554/eLife.64972.

[39] Aly M, Turk-Browne NB. Attention stabilizes representations in the human hippocampus. Cereb Cortex 2016;26:783–96. 10.1093/cercor/bhv041.

[40] Aminoff E, Gronau N, Bar M. The parahippocampal cortex mediates spatial and nonspatial associations. Cereb Cortex 2007;17:1493–503. 10.1093/cercor/bhl078.

[41] Ranganath C. A unified framework for the functional organization of the medial temporal lobes and the phenomenology of episodic memory. Hippocampus 2010;20:1263–90. 10.1002/hipo.20852.

[42] Kornysheva K, Bush D, Meyer SS, Sadnicka A, Barnes G, Burgess N. Neural competitive queuing of ordinal structure underlies skilled sequential action. Neuron 2019;101:1166–1180.e3. 10.1016/j.neuron.2019.01.018.

[43] Yokoi A, Arbuckle SA, Diedrichsen J. The role of human primary motor cortex in the production of skilled finger sequences. J Neurosci 2018;38:1430–42. 10.1523/jneurosci.2798-17.2017.

[44] Berlot E, Popp NJ, Diedrichsen J. A critical re-evaluation of fMRI signatures of motor sequence learning. Elife 2020;9. 10.7554/eLife.55241.

[45] Yokoi A, Diedrichsen J. Neural organization of hierarchical motor sequence representations in the human neocortex. Neuron 2019;103:1178–1190.e7. 10.1016/j.neuron.2019.06.017.

[46] Faul F, Erdfelder E, Lang A-G, Buchner A. G*Power 3: a flexible statistical power analysis program for the social, behavioral, and biomedical sciences. Behav Res Methods 2007;39:175–91. 10.3758/bf03193146.

[47] Oldfield RC. The assessment and analysis of handedness: the Edinburgh inventory. Neuropsychologia 1971;9:97–113. 10.1016/0028-3932(71)90067-4.

[48] Beck AT, Epstein N, Brown G, Steer RA. An inventory for measuring clinical anxiety: psychometric properties. J Consult Clin Psychol 1988;56:893–7. 10.1037//0022-006x.56.6.893.

[49] Beck AT, Ward CH, Mendelson M, Mock J, Erbaugh J. An inventory for measuring depression. Arch Gen Psychiatry 1961;4:561–71. 10.1001/archpsyc.1961.01710120031004.

[50] Buysse DJ, Reynolds CF 3rd, Monk TH, Berman SR, Kupfer DJ. The Pittsburgh Sleep Quality Index: a new instrument for psychiatric practice and research. Psychiatry Res 1989;28:193–213. 10.1016/0165-1781(89)90047-4.

[51] Johns MW. A new method for measuring daytime sleepiness: the Epworth sleepiness scale. Sleep 1991;14:540–5. 10.1093/sleep/14.6.540.

[52] Horne JA, Ostberg O. A self-assessment questionnaire to determine morningness-eveningness in human circadian rhythms. Int J Chronobiol 1976;4:97–110.

[53] Kleiner M, Brainard D, Pelli D. 2007.

[54] Brown RM, Robertson EM. Off-line processing: reciprocal interactions between declarative and procedural memories. J Neurosci 2007;27:10468–75. 10.1523/JNEUROSCI.2799-07.2007.

[55] Ellis BW, Johns MW, Lancaster R, Raptopoulos P, Angelopoulos N, Priest RG. The St. Mary’s Hospital sleep questionnaire: a study of reliability. Sleep 1981;4:93–7. 10.1093/sleep/4.1.93.

[56] MacLean AW, Fekken GC, Saskin P, Knowles JB. Psychometric evaluation of the Stanford Sleepiness Scale. J Sleep Res 1992;1:35–9. 10.1111/j.1365-2869.1992.tb00006.x.

[57] Pan SC, Rickard TC. Sleep and motor learning: Is there room for consolidation? Psychol Bull 2015;141:812–34. 10.1037/bul0000009.

[58] Tompary A, Davachi L. Editorial note to: Consolidation promotes the emergence of representational overlap in the hippocampus and medial prefrontal cortex. Neuron 2020;105:198. 10.1016/j.neuron.2019.12.021.

[59] Fan L, Li H, Zhuo J, Zhang Y, Wang J, Chen L, et al. The human brainnetome atlas: A new brain atlas based on connectional architecture. Cereb Cortex 2016;26:3508–26. 10.1093/cercor/bhw157.

[60] Liuzzi AG, Bruffaerts R, Vandenberghe R. The medial temporal written word processing system. Cortex 2019;119:287–300. 10.1016/j.cortex.2019.05.002.

