## Supplementary Material for "The capacity of the medial temporal lobe to represent memory items in their ordinal position in a sequence is domain-general"

**This PDF file includes:**

Supporting text

Figures S1 to S2

Tables S1 to S6

### Supplementary results

#### 1. Assessment of baseline performance

We tested whether general motor execution, assessed with the random SRTT, differed at baseline between the 2 learning sessions on day 1 (see Fig. 1A in the main text). Results show that performance speed (i.e., mean response time) increased across blocks of the random SRTT (block effect: $F(3,87)=7.71$, $\eta_p^2=0.21$, $p<0.001$) but this effect did not differ between sessions (session by block effect: $F(3,87)=0.64$, $\eta_p^2=0.02$, $p=0.59$; session effect $F(1,29)=0.02$, $\eta_p^2=0.001$, $p=0.89$; see upper panel of Fig. 2 in the main text). Performance accuracy (i.e., % correct responses) remained stable across blocks of the random SRTT (session by block effect: $F(3,87)=0.39$, $\eta_p^2=0.01$, $p=0.76$; block effect: $F(3,87)=1.07$, $\eta_p^2=0.04$, $p=0.37$; see bottom panel of Fig. 2 in the main text) but was better during the object task session (session effect; $F(1,29)=5.9$, $\eta_p^2=0.17$, $p=0.02$).

#### 2. Session order effects on performance

Even though condition order was counterbalanced across participants on experimental days 1 (learning) and 2 (retest), we tested whether order influenced performance on both the random and sequence SRTT. Results of the ANOVAs using session order, irrespective of task condition and block as within-subject factors are presented below.

Day 1 data

For the random SRTT, performance speed increased across blocks (block effect: $F(3,87)=7.71$, $\eta_p^2=0.21$, $p<0.001$) and was overall better in session 2 than in session1 (session by block effect: $F(3,87)=8.86$, $\eta_p^2=0.23$, $p<0.001$; session effect $F(1,29)=87.23$ $\eta_p^2=0.75$, $p<0.01$). In contrast, accuracy remained stable across blocks of the random SRTT (block effect: $F(3,87)=1.07$, $\eta_p^2=0.04$, $p=0.37$) and similar across sessions (session effect; $F(1,29)=0.41$, $\eta_p^2=0.01$, $p=0.53$; session by block effect: $F(3,87)=1$, $\eta_p^2=0.03$, $p=0.4$).

For the sequential SRTT, performance speed improved during training in both sessions (block effect; training: $F(19,551)=38.98$, $\eta_p^2=0.57$, $p<0.001$; test: $F(3,87)=3.78$, $\eta_p^2=0.12$, $p=0.01$) and the block by session interaction was significant such that block-to-block changes in performance were steeper in the early blocks of session 2 as compared to session 1 (condition by block effect; training: $F(19,551)=4.92$, $\eta_p^2= 0.15$, $p<0.001$; note that this effect was no longer observed during test: $F(3,87)=0.33$, $\eta_p^2=0.01$, $p=0.8$). However, overall performance speed did not differ between sessions (session effect; training: $F(1,29)=0.95$, $\eta_p^2=0.03$, $p=0.34$; test: $F(1,29)=0.58$, $\eta_p^2= 0.02$, $p=0.45$). Performance accuracy remained similar across sessions (training: session by block effect: $F(19,551)=1.64$, $\eta_p^2=0.05$, $p=0.04$; block effect: $F(19,551)=1.43$, $\eta_p^2=0.05$, $p=0.11$; session effect: $F(1,29)=0.02$, $\eta_p^2=0.001$, $p=0.89$; test: session by block effect: $F(3,87)=1.96$, $\eta_p^2=0.06$, $p=0.13$; block effect $F(3,87)=0.72$, $\eta_p^2=0.02$, $p=0.55$; session effect: $F(1,29)=0.19$, $\eta_p^2=0.01$, $p=0.66$).

Day 2 data

During the retest session on day 2, performance speed improved across practice blocks while accuracy remained stable (block effect; speed: $F(3,87)=9.07$, $\eta_p^2=0.24$, $p<0.001$; accuracy: $F(3,87)=1.70$, $\eta_p^2=0.06$, $p=0.17$). Performance speed was faster for the task that was retested second, while accuracy was similar between sessions (order effect; speed: $F(1,29)=6.65$; $\eta_p^2=0.19$, $p=0.02$; accuracy: $F(1,29)=0.05$, $\eta_p^2=0.002$, $p=0.83$)

Overall, these results suggest that on day 1, participants tended to be faster at the beginning of session 2 as compared to session 1 (random task and early blocks of sequence tasks) which suggests skill transfer between sessions. An order effect was also observed on day 2 during retest outside the scanner. However, it is unlikely that these order effects influenced our results as the order of the conditions was counterbalanced across participants (15 participants started with the motor condition and 15 participants started with the object condition).

#### 3. Brain- behavioral correlation

Since performance differed between conditions during initial learning, we examined whether the level of performance reached at the end of the training session on day 1 (average reaction time in post-training test blocks) was correlated with neural pattern similarity (delta similarity for object-position and finger-position coding) examined the next day. We did not observe any correlation between performance on the motor sequence task and the finger-position coding effect in any of the motor ROIs (M1: r=-0.38, pcorr=0.2; PMC: r=-0.45, pcorr=0.06; HC: r=-0.32, pcorr=0.42) or between performance on the object sequence task and the object-position coding effect in any of the object ROIs (PHC: r=-0.27, pcorr=0.73; PER: r=-0.28, pcorr=0.68; HC: r=-0.2, pcorr=1). Altogether, these results indicate that the level of performance reached at the end of training on day 1 was not related to the multivoxel activation patterns examined the next day.

#### Supplementary Figures

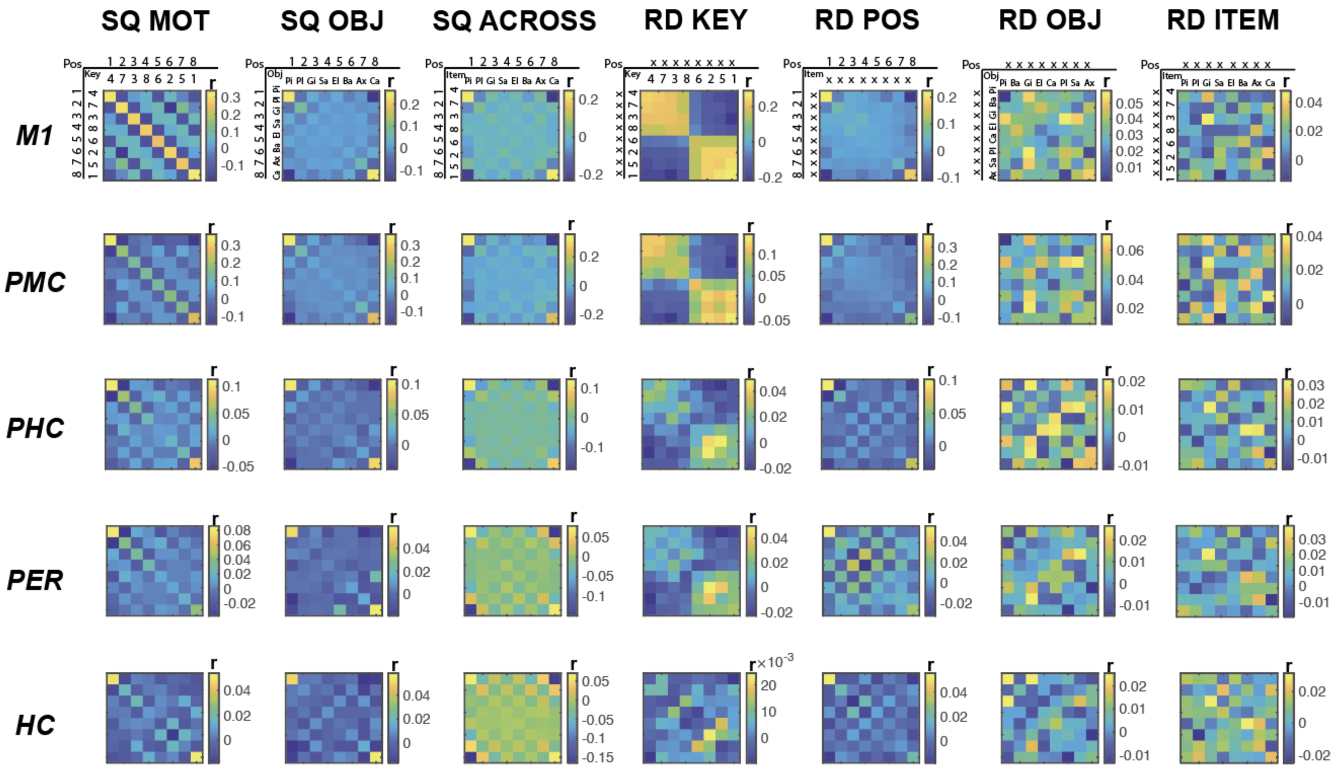


***Figure S1*. *Similarity Matrices.*** *Group average neural similarity matrices for all the ROIs. Pattern similarity was computed across repetitions of the motor sequence to assess* ***finger-position*** *coding in the sequence condition (**SQ MOT matrix**), across repetitions of the object sequence to assess* ***object-position*** *coding in the sequence condition (**SQ OBJ matrix**), between pairs of the individual objects from the object sequence condition and individual fingers from the motor sequence condition to assess* ***item-position*** *coding across sequences from different domains (**SQ ACROSS matrix**), as well as across repetitions of the random patterns to quantify* ***finger/key*** *(**RD KEY matrix**),* ***position*** *(**RD POS matrix**),* ***object*** *(**RD OBJ matrix**) and* ***item*** *(**RD ITEM**) coding in the random condition. Color bar represents mean similarity (r). In the POS rows/columns, numbers represent the temporal position in a sequence. In the KEY rows/columns, numbers represent fingers. In the OBJ rows/columns, the first 2 letters of each object are presented. (X) represents a random position or key/object in the item and position matrices, respectively.*

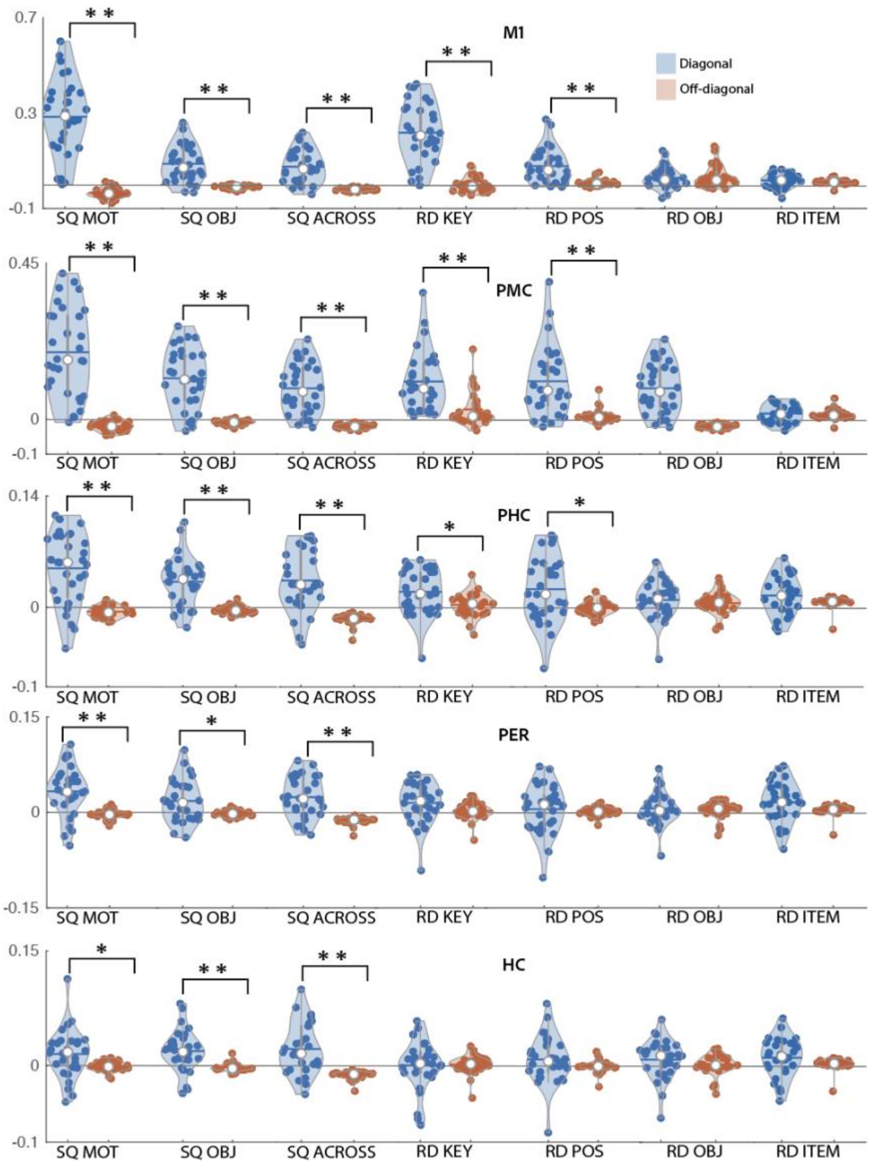


***Figure S2*. Mean pattern similarity for diagonal (blue) and off-diagonal (red) cells as a function of matrices and ROIs**. *Results indicate that all ROIs (primary motor cortex (M1), premotor cortex (PMC), parahippocampus (PHC), perirhinal (PER), hippocampus (HC)) show evidence of finger-position, object-position and item-position coding in the sequence conditions. M1, PMC and PHC show evidence of finger and position coding during random practice whereas HC and PER do not. None of the ROIs showed evidence of object or item coding. Asterisks indicate significant differences between diagonal and off-diagonal (one sided paired sample t-test; Bonferroni corrected* $*p_{corr}<.05$ *and* $** \; p_{corr}<.005$*). Colored circles represent individual data, jittered in arbitrary distances on the x-axis to increase perceptibility. Horizontal lines represent means and white circles represent medians. The shape of the violin [1] depicts the kernel density estimate of the data. Note that as in earlier research [16], Y axis scales are different between ROIs to accommodate for differences in signal-to-noise ratio (and therefore in effect sizes) between ROIs.*

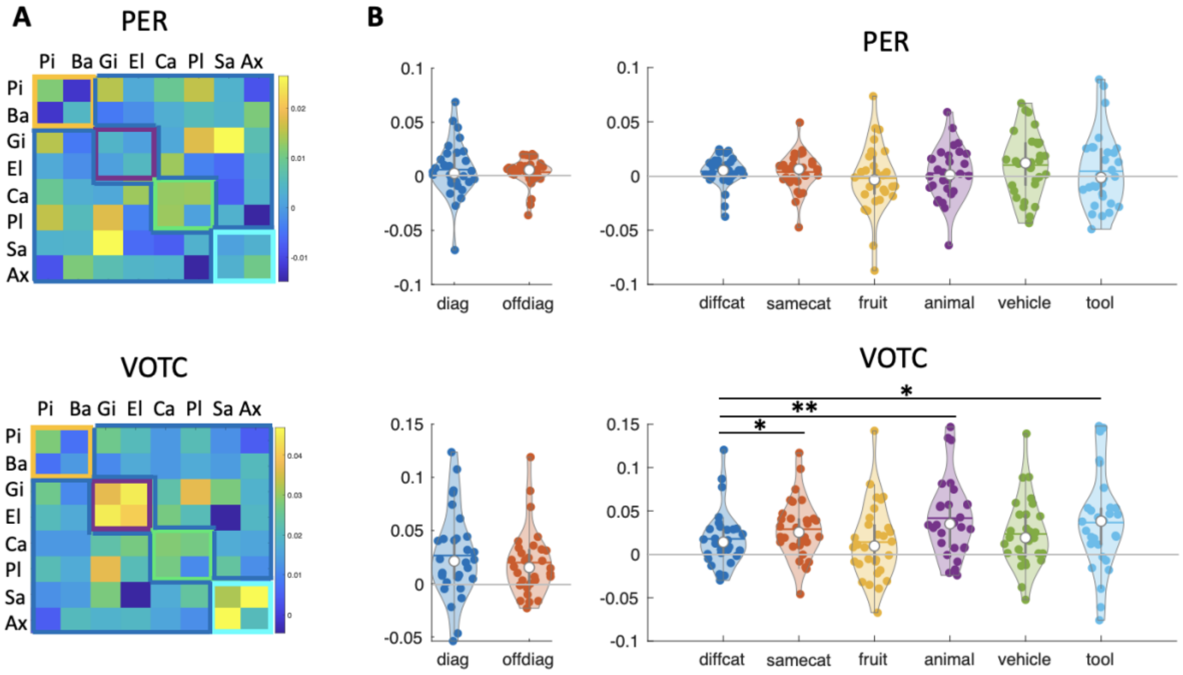


***Figure S3*. Object category analysis** *A. Group average neural similarity matrices for objects using random data in the perirhinal cortex (PER) and the ventro-occipo-temporal cortex (VOTC) to assess object/object category coding. Colored squares along the diagonal represent correlations between object within the same category, i.e., fruit (yellow outline), animal (purple outline), vehicle (green outline) and tool (blue outline) categories. In the rows/columns, the first 2 letters of each object are presented. B. Left panel. Mean pattern similarity for diagonal (correlation between repetitions of the same object) and off-diagonal (correlation between repetition of different objects) cells. In line with the results presented in the main text, these analyses indicate that neither PER nor VOTC showed evidence of object coding. Right panel. Mean pattern similarity between different object categories (diffcat, see dark blue frame in matrices presented in panel A), between same object categories (samecat, i.e. across all cells highlighted with the four colored frames along the diagonal in matrices presented in Panel A) and within each of the object categories (fruit, animal, vehicle, tool). Results showed a main effect of category (same vs. different) in the VOTC (t(29)=2.78, d=0.51, p=0.01) which confirms the well-known object category coding in this region (see discussion in main text). This effect was particularly pronounced for the animal (animal category vs. different category, t(29)=4.25, d=0.78, p<0.002) and tool (tool category vs. different category, t(29)=2.25, d=0.41, p=0.04) categories. No such effects were observed in the PER. Asterisks indicate significant differences (one sided paired sample t-test; Bonferroni corrected *$p_r$<.05 and ** $p_r$<.005). Colored circles represent individual data, jittered in arbitrary distances on the x-axis to increase perceptibility. Horizontal lines represent means and white circles represent medians. The shape of the violin [22] depicts the kernel density estimate of the data.*

#### Supplementary Tables

| ROI | T-stat | DF | pcorr |
|---|---|---|---|
| M1 | 2.248 | 29 | 0.08 |
| PMC | 3.036 | 29 | **0.02** |
| PHC | -0.156 | 29 | 2.2 |
| PER | -1.402 | 29 | 0.43 |
| HC | -0.419 | 29 | 1 |

***Table S1. Control analyses on 6x6 matrices after removing boundary positions.*** *One-tailed one sample-test results for the comparison of delta similarity (mean diagonal cells vs. mean off-diagonal cells) against zero for position matrices after edge removal (6x6 matrices). After removing boundary positions, position coding only remained significant in the premotor cortex while such coding was no longer significant in all other ROIs. P values are Bonferroni-corrected.*

| **M1** | | | | **Step1** | | | | **Step2** | |
|---|---|---|---|---|---|---|---|---|---|
| **stepwise** | **predictors** | **B** | **beta** | **t** | **pcorr** | **B** | **beta** | **t** | **pcorr** |
| | key | 1.252 | 0.833 | 7.977 | **<0.005** | 0.898 | 0.598 | 6.499 | **<0.005** |
| | pos | | | | | 1.184 | 0.444 | 4.828 | **<0.005** |
| | **Model** | **R2** | **F** | **△R2** | **pcorr** | **R2** | **F** | **△R2** | **pcorr** |
| | | 0.694 | 63.638 | 0.694 | **<0.005** | 0.836 | 68.821 | 0.142 | **<0.005** |
| **PMC** | | | | **Step1** | | | | **Step2** | |
| **stepwise** | **predictors** | **B** | **beta** | **t** | **pcorr** | **B** | **beta** | **t** | **pcorr** |
| | key | | | | | 0.773 | 0.333 | 4.081 | **<0.005** |
| | pos | 1.362 | 0.87 | 9.322 | **<0.005** | 1.152 | 0.736 | 9.018 | **<0.005** |
| | **Model** | **R2** | **F** | **△R2** | **pcorr** | **R2** | **F** | **△R2** | **pcorr** |
| | | 0.756 | 86.904 | 0.756 | **<0.005** | 0.849 | 76.079 | 0.093 | **<0.005** |
| **HC** | | | | **Step1** | | | | **Step2** | |
| **stepwise** | **predictors** | **B** | **beta** | **t** | **pcorr** | **B** | **beta** | **t** | **pcorr** |
| | key | | | | | 0.065 | 0.052 | 0.335 | 3.7 |
| | pos | 0.624 | 0.602 | 3.994 | **<0.005** | 0.632 | 0.611 | 3.934 | **<0.005** |
| | **Model** | **R2** | **F** | **△R2** | **pcorr** | **R2** | **F** | **△R2** | **pcorr** |
| | | 0.363 | 15.954 | 0.363 | **<0.005** | 0.366 | 7.78 | 0.003 | 3.7 |
| **PER** | | | | **Step1** | | | | **Step2** | |
| **stepwise** | **predictors** | **B** | **beta** | **t** | **pcorr** | **B** | **beta** | **t** | **pcorr** |
| | key | 0.596 | 0.449 | 2.658 | 0.065 | 0.578 | 0.435 | 2.686 | 0.06 |
| | pos | | | | | 0.324 | 0.304 | 1.878 | 0.355 |
| | **Model** | **R2** | **F** | **△R2** | **pcorr** | **R2** | **F** | **△R2** | **pcorr** |
| | | 0.201 | 7.062 | 0.201 | 0.065 | 0.294 | 5.614 | 0.92 | 0.355 |
| **PHC** | | | | **Step1** | | | | **Step2** | |
| **stepwise** | **predictors** | **B** | **beta** | **t** | **pcorr** | **B** | **beta** | **t** | **pcorr** |
| | key | | | | | 0.643 | 0.368 | 2.561 | 0.08 |
| | pos | 0.63 | 0.562 | 3.599 | **0.005** | 0.583 | 0.521 | 3.626 | **0.005** |
| | **Model** | **R2** | **F** | **△R2** | **pcorr** | **R2** | **F** | **△R2** | **pcorr** |
| | | 0.316 | 12.953 | 0.316 | **0.005** | 0.45 | 11.042 | 0.134 | 0.08 |

***Table S2. Stepwise linear regression results for motor sequence learning for all ROIs.*** *Dependent variable: delta similarity (i.e., difference between mean diagonal and mean off-diagonal cells) in the motor sequence matrix. Predictors: delta similarity in the random key and/or position matrix. An F-test was used to assess whether the change in explained variance (△R 2) from the prior step is significant. Step 1, df1=1, df2=28; Step 2, df1=2, df2=27. P-values are Bonferroni corrected.*

| PER | | Step1 | | | | Step2 | | | |
|---|---|---|---|---|---|---|---|---|---|
| **stepwise** | **predictors** | **B** | **beta** | **t** | **pcorr** | **B** | **beta** | **t** | **pcorr** |
| | object | | | | | 0.454 | 0.306 | 1.826 | 0.395 |
| | pos | 0.346 | 0.385 | 2.21 | 0.175 | 0.333 | 0.371 | 2.213 | 0.18 |
| | **Model** | **R2** | **F** | **△R2** | **pcorr** | **R2** | **F** | **△R2** | **pcorr** |
| | | 0.149 | 4.886 | 0.149 | 0.175 | 0.242 | 4.315 | 0.94 | 0.395 |
| **PHC** | | **Step1** | | | | **Step2** | | | |
| **stepwise** | **predictors** | **B** | **beta** | **t** | **pcorr** | **B** | **beta** | **t** | **pcorr** |
| | object pos | | | | | 0.63 | 0.388 | 2.962 | **0.03** |
| | | 0.491 | 0.629 | 4.28 | **<0.005** | 0.455 | 0.583 | 4.455 | **<0.005** |
| | **Model** | **R2** | **F** | **△R2** | **pcorr** | **R2** | **F** | **△R2** | **pcorr** |
| | | 0.395 | 18.317 | 0.393 | **<0.005** | 0.544 | 16.09 | 0.148 | **0.03** |
| **HC** | | **Step1** | | | | **Step2** | | | |
| **stepwise** | **predictors** | **B** | **beta** | **t** | **pcorr** | **B** | **beta** | **t** | **pcorr** |
| | object | | | | | 0.212 | 0.171 | 1.01 | 1 |
| | pos | 0.408 | 0.449 | 2.658 | 0.065 | 0.391 | 0.431 | 2.537 | 0.085 |
| | **Model** | **R2** | **F** | **△R2** | **pcorr** | **R2** | **F** | **△R2** | **pcorr** |
| | | 0.201 | 7.063 | 0.201 | 0.065 | 0.23 | 4.044 | 0.029 | 1 |
| **M1** | | **Step1** | | | | **Step2** | | | |
| **stepwise** | **predictors** | **B** | **beta** | **t** | **pcorr** | **B** | **beta** | **t** | **pcorr** |
| | object | | | | | -0.34 | -0.121 | -1.255 | 1 |
| | pos | 1.077 | 0.875 | 9.407 | **<0.005** | 1.032 | 0.839 | 8.691 | **<0.005** |
| | **Model** | **R2** | **F** | **△R2** | **pcorr** | **R2** | **F** | **△R2** | **pcorr** |
| | | 0.766 | 88.497 | 0.766 | **<0.005** | 0.78 | 45.976 | 0.013 | 1.105 |
| **PMC** | | **Step1** | | | | **Step2** | | | |
| **stepwise** | **predictors** | **B** | **beta** | **t** | **pcorr** | **B** | **beta** | **t** | **pcorr** |
| | object | | | | | 0.08 | 0.026 | 0.268 | 0.396 |
| | pos | 0.879 | 0.876 | 9.63 | **<0.005** | 0.887 | 0.884 | 9.094 | **<0.005** |
| | **Model** | **R2** | **F** | **△R2** | **pcorr** | **R2** | **F** | **△R2** | **pcorr** |
| | | 0.768 | 92.745 | 0.768 | **<0.005** | 0.769 | 44.871 | 0.001 | 0.396 |

***Table S3. Stepwise linear regression results for object sequence learning for all ROIs.*** *Dependent variable: delta similarity (i.e., difference between mean diagonal and mean off-diagonal cells) in the object sequence matrix. Predictors: delta similarity in the random object and/or position matrix. An F-test was used to assess whether the change in explained variance (△R 2) from the prior step is significant. Step 1, df1=1, df2=28; Step 2, df1=2, df2=27. P-values are Bonferroni corrected.*

| **M1** | | **Step1** | | | | **Step2** | | | |
|---|---|---|---|---|---|---|---|---|---|
| **step wise** | **predictors** | **B** | **beta** | **t** | **pcorr** | **B** | **beta** | **t** | **pcorr** |
| | item | | | | | 0.105 | 0.049 | 0.397 | 1 |
| | pos | 0.887 | 0.812 | 7.369 | **<0.005** | 0.91 | 0.833 | 6.728 | **<0.005** |
| | **Model** | **$R^2$** | **F** | **$\triangle R^2$** | **pcorr** | **$R^2$** | **F** | **$\triangle R^2$** | **pcorr** |
| | | 0.66 | 54.305 | 0.66 | **<0.005** | 0.662 | 26.415 | 0.002 | 1 |
| **PMC** | | **Step1** | | | | **Step2** | | | |
| **step wise** | **predictors** | **B** | **beta** | **t** | **pcorr** | **B** | **beta** | **t** | **pcorr** |
| | item | | | | | 0.206 | 0.081 | 0.771 | 1 |
| | pos | 0.723 | 0.871 | 9.392 | **<0.005** | 0.754 | 0.908 | 8.64 | **<0.005** |
| | **Model** | **$R^2$** | **F** | **$\triangle R^2$** | **pcorr** | **$R^2$** | **F** | **$\triangle R^2$** | **pcorr** |
| | | 0.759 | 88.215 | 0.759 | **<0.005** | 0.764 | 43.765 | 0.005 | 1 |
| **PER** | | **Step1** | | | | **Step2** | | | |
| **step wise** | **predictors** | **B** | **beta** | **t** | **pcorr** | **B** | **beta** | **t** | **pcorr** |
| | item | | | | | -0.077 | -0.078 | -0.43 | 1 |
| | pos | 0.271 | 0.34 | 1.916 | 0.33 | 0.265 | 0.332 | 1.833 | 0.39 |
| | **Model** | **$R^2$** | **F** | **$\triangle R^2$** | **pcorr** | **$R^2$** | **F** | **$\triangle R^2$** | **pcorr** |
| | | 0.116 | 3.669 | 0.116 | 0.33 | 0.122 | 1.874 | 0.006 | 3.35 |
| **PHC** | | **Step1** | | | | **Step2** | | | |
| **step wise** | **predictors** | **B** | **beta** | **t** | **pcorr** | **B** | **beta** | **t** | **pcorr** |
| | item | | | | | -0.072 | -0.056 | -0.345 | 1 |
| | pos | 0.511 | 0.581 | 3.775 | **<0.005** | 0.498 | 0.567 | 3.505 | **0.01** |
| | Model | $R^2$ | F | $\triangle R^2$ | **pcorr** | $R^2$ | F | $\triangle R^2$ | **pcorr** |
| | | 0.337 | 14.249 | 0.337 | **<0.005** | 0.34 | 6.96 | 0.003 | 3.665 |
| **HC** | | **Step1** | | | | **Step2** | | | |
| **step wise** | **predictors** | **B** | **beta** | **t** | **pcorr** | **B** | **beta** | **t** | **pcorr** |
| | item | | | | | -0.109 | -0.103 | -0.603 | 1 |
| | pos | 0.436 | 0.446 | 2.635 | 0.07 | 0.443 | 0.453 | 2.641 | 0.07 |
| | **Model** | **$R^2$** | **F** | **$\triangle R^2$** | **pcorr** | **$R^2$** | **F** | **$\triangle R^2$** | **pcorr** |
| | | 0.199 | 6.946 | 0.199 | 0.07 | 0.209 | 3.575 | 0.011 | 1 |

***Table S4. Stepwise linear regression results for sequence learning across domains for all ROIs.*** *Dependent variable: delta similarity (i.e., difference between mean diagonal and mean off-diagonal cells) in the item-position matrix. Predictors: delta similarity in the random item and/or position matrix. An F-test was used to assess whether the change in explained variance ($\triangle R^2$) from the prior step is significant. Step 1, df1=1, df2=28; Step 2, df1=2, df2=27. P-values are Bonferroni corrected.*

| **N** | 30 (18 Females) |
|---|---|
| Age (yrs) | 23.3 ± 5.21 |
| Edinburgh Handedness [47] (>40) | 77.67 ± 17.85 |
| Epworth Sleepiness Scale [51] (<10) | 4.33 ± 3.3 |
| Beck Depression Scale [49] (<17) | 2.97 ± 3.9 |
| Beck Anxiety Scale [48] (<17) | 2.93 ± 3.45 |
| PSQI [50] (<8) | 2.87 ± 2 |
| Chronoscore (CRQ, [52]) (30<n<70) | 55.87 ± 9.79 |
| **Sleep duration in hrs (Sleep diary)** | |
| Night 1 | 8.32 ± 1.51 |
| Night 2 | 8.6 ± 1.48 |
| Night 3 | 8 ± 1.14 |
| Night 4[a] | 8.77 ± 1.84 |
| One-way rmANOVA results | F(3,87)=2.52; *P*-value = 0.06 |
| **St. Mary's questionnaire** | |
| Duration in hrs (Night 3) | 7.73 ± 0.82 |
| Duration in hrs (Night 4) | 8.12 ± 1.18 |
| Paired sample T-test | t(29)=-1.85; P-value = 0.07 |
| Quality (Night 3)[b] | 4.83 ± 0.83 |
| Quality (Night 4) | 4.83 ± 0.95 |
| Paired sample T-test | t(29)=-0.0; P-value = 1 |
| **Between session intervals (hrs)** | |
| Session1-Session2 | 4.78 ± 1.05 |
| Session2-Session3 | 23.7 ± 1.6 |
| **Psychomotor Vigilance Task[c] (s)** | |
| Session 1 | 0.32 ± 0.04 |
| Session 2 | 0.32 ± 0.04 |
| Session 3 | 0.31 ± 0.04 |
| One-way rmANOVA results | F(2,58)=0.89; *P*-value = 0.42 |
| **Stanford sleepiness score[d]** | |
| Session 1 | 1.77 ± 0.57 |
| Session 3 | 1.7 ± 0.7 |
| Paired sample T-test | t(29)=0.4; P-value = 0.7 |

***Table S5*. *Participant demographics, sleep and vigilance characteristics.*** *Notes. Values are means and standard deviations (unless noted otherwise). Cutoff values for exclusion in parentheses. PSQI = Pittsburgh Sleep Quality Index. CRQ = Circadian Rhythm Questionnaire. [a] Determined in combination between sleep diary and wrist actigraphy recordings on night 4 (between the two experimental sessions). [b]1=very badly, 6 very well. [c] Median of reaction times. [d]1=more alert, 7=less alert.*

| ROI | Mean | Range |
|---|---|---|
| M1 | 2628 | 2226-3034 |
| PMC | 2428 | 2023-3026 |
| PHC | 989 | 808-1124 |
| PER | 542 | 255-713 |
| HC | 581 | 442-728 |

***Table S6*. *ROI size* *(number of voxels).***
